# Inferring sources of suboptimality in perceptual decision making using a causal inference task

**DOI:** 10.1101/2022.04.28.489925

**Authors:** Sabyasachi Shivkumar, Madeline S. Cappelloni, Ross K. Maddox, Ralf M. Haefner

## Abstract

Perceptual decision-making has been extensively modeled using the ideal observer framework. However, a range of deviations from optimality demand an extension of this framework to characterize the different sources of suboptimality. Prior work has mostly formalized these sources by adding biases and variability in the context of specific process models but are hard to generalize to more complex tasks. Here, we formalize suboptimalities as part of the brain’s probabilistic model of the task. Data from a traditional binary discrimination task cannot separate between different kinds of biases, or between sensory noise and approximate computations. We showed that this was possible using a recently developed causal inference task in which observers discriminated auditory cues in the presence of choice-uninformative visual cues. An extension of the task with different stimulus durations provided evidence for an increase in the precision of the computations with stimulus duration, separate from a decrease in observation noise.

## 1 Introduction

Much of our knowledge about how the brain converts sensory observations into percepts, and percepts into decisions, has been gained in the context of binary discrimination tasks. In such tasks, human behavior has often been found to be close to optimal under certain sensory noise distributions (Swets et al., 1961; Ernst and Banks, 2002). Normative modeling starts by specifying the experimenter’s probabilistic model that links observations to correct choices. It assumes that the brain has learned and uses this model to infer correct responses from its observations. These models are mathematical descriptions of how an optimal decision making agent (ideal observer) would make their decisions. Approaching complex systems like the brain through the lens of the ideal observer has provided us with guiding principles (Geisler, 2011). However, there is increasing evidence for suboptimalities in human behavior (reviewed in Rahnev and Denison (2018)) calling into question the utility of a normative approach (Bowers and Davis, 2012; Gardner, 2019). Instead of abandoning the ideal observer model completely, it has been suggested to use it as a starting point (Icard, 2018). To construct better models of human behavior it can be extended by incorporating increasingly realistic assumptions about its components (Rahnev and Denison, 2018).

There are two principal approaches to this process. The first approach starts with the generative model for the task. Suboptimalities arise from deviations in the brain’s internal model, and the fact that the inference computations are performed approximately instead of exactly. For example, systematic biases in observer responses have been modeled as arising due to observers using priors learned for natural behavior that are not optimal in laboratory settings (Stocker and Simoncelli, 2006; Odegaard et al., 2015; Ma, 2019). Similarly, approximations in the inference process can lead to substantial deviations from optimal behavior and have been shown to be a important source of suboptimality in decisions (Beck et al., 2012; Wyart and Koechlin, 2016; Drugowitsch et al., 2016). One way in which approximations in the inference computations have been quantified is by using a sampling-based approximation, where the number of samples quantifies the degree of approximation. Prior studies have found that observers are best described by few samples, corresponding to coarse approximations (Vul et al., 2014; Sanborn and Chater, 2016; Wozny et al., 2010).

The second approach starts with a process model that inverts the above generative model. This model optimally converts observations into responses (as defined for the ideal observer) and suboptimalities can be added to its components in the form of noise or bias. This approach was followed in recent attempts to dissociate between sensory and computational sources of suboptimality (Drugowitsch et al., 2016), and different sources of biases (Linares et al., 2019). Most commonly, this approach uses the signal detection theory framework where a decision is made by comparing the observation against a criterion (Gold and Shadlen, 2007; Green et al., 1966). Response biases can then be modeled as an incorrect placement of the criterion, and response variability as an inability to maintain a stable criterion (see Rahnev and Denison (2018) for a detailed review). Formalizing suboptimalities in a process model introduces the challenge that complicated probabilistic models (for more complex or naturalistic tasks) do not allow for implementation-agnostic models to which variability or a bias can be applied as commonly done. Instead they require a commitment to how the inference computations are being implemented despite the fact that this implementation in the brain is still unknown (Pouget et al., 2013; Fiser et al., 2010). This problem is compounded by the fact that it is unclear to what degree different sources of suboptimality can be dissociated given empirical data from simple tasks (also a challenge in the first approach), driving the need for more complex tasks. For instance, (Linares et al., 2019) combined data from two task conditions to be able to dissociate perceptual biases from category biases. In order to dissociate sensory noise from approximate inference, (Lengyel et al., 2015) designed a dual-report estimation/confidence judgement task, and (Drugowitsch et al., 2016) designed a task requiring the accumulation of evidence across a variable number of pieces of evidence. Therefore separating sources of suboptimalities has required a need for using more complex tasks.

Here, we followed the first approach. We derived a formalization of decision making in binary discrimination tasks under common Gaussian assumptions and showed that data from classic discrimination tasks cannot distinguish between perceptual and categorical biases, nor between sensory noise and approximate computations. In order to distinguish between these different sources of suboptimality, we applied our formalization to a hierarchical causal inference model of audio-visual integration. Analyzing previously collected data (Cappelloni et al., 2019), we showed that data from this task dissociates all four sources of suboptimal inference described above. We also applied our model to an extension of the task with variable duration (Cappelloni et al., 2020) and showed that both the computational approximation becomes coarser, and the observation noise becomes larger, as the duration of the stimulus becomes shorter.

## 2 Results

Our Results section is organized as follows: we first formalize the different sources of suboptimalities in the context of classic binary discrimination tasks and demonstrate how different sources of bias and variability cannot be dissociated using data from classic discrimination tasks. We next apply the same formalization to a recent task which does allow for such a dissociation. Finally, we show that the improvement in behavioral performance with increasing stimulus duration is the result of *both* less sensory noise *and* more precise computations.

### 2.1 Sources of approximate decision-making in a binary discrimination task

In traditional binary discrimination tasks where the observer has to compare a cue against a reference boundary, the observer’s responses can be summarized using a psychometric curve (observer reports measured as a function of cue position). The psychometric curve is commonly characterized by bias(Figure 1A) and a threshold (Figure 1B). The bias can be equivalently measured (Figure 1A) as the proportion of ‘right’ responses for cue position at the midline or the point of subjective equality (PSE) which is the cue position for which the observer reports ‘right’ and ‘left’ with equal probability. The threshold (Figure 1B) is proportional to the inverse slope around the PSE.

**Figure 1:**
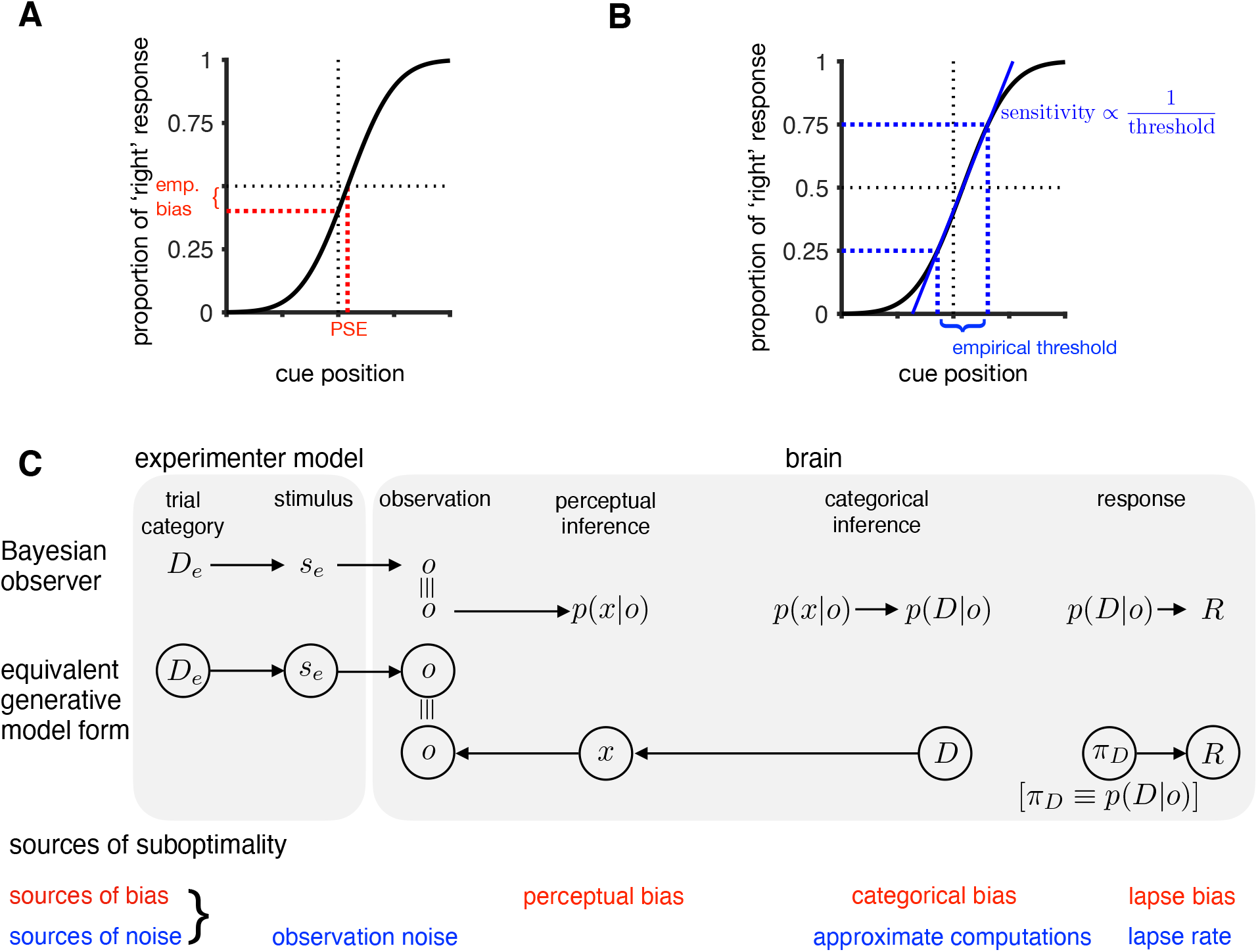
Modeling suboptimalities in perceptual decision making in a binary discrimination task: **(A,B)** Predicted psychometric curve for a Bayesian observer. Responses can be summarized using the empirical bias (or equivalently the point of subjective equality, i.e. PSE) as shown in A and the empirical threshold (or equivalently the slope around PSE, i.e. sensitivity) as shown in B. **(C) Top row:** Stages of perceptual decision making: In a binary discrimination task, where the goal is to discriminate which side of a reference the presented cue came from, the Bayesian observer first infers the belief about the cue position (*x*). It does so by combining the likelihood of the noisy observation (*o*) with its prior belief about the cue position. The noisy observation differs from the experimenter defined cue position (*s_e_*) due to external and internal sources of noise. The ideal observer then infers the belief about the trial category (*D*) which is then converted to a response in the decision making stage where the response is chosen according to Bayesian decision theory; **Middle row:** Equivalent generative model formalization of the decision making process. **Bottom row:** Suboptimalities in the decision making process. The observer response could differ from the experimenter chosen correct trial category (*D_E_*) due to biases and noise/approximations associated with the different parts of each model.

We formalize optimal decision-making by modeling it as Bayesian inference. We assume that the brain has learned an approximation to the generative model of the task, and that it inverts this model for inference and decision-making (Figure 1C). On every trial the experimenter chooses a ‘correct’ trial category (*D_e_*) and a stimulus, i.e. cue (*s_e_*). For concreteness, we will use a position discrimination task in line with the second part of the paper as an example but the formalization here holds in general. The observer observes a noisy version of the cue position that deviates from the experimenter defined value due to sources of noise that may be external or internal from the brain’s perspective. The ideal observer computes the posterior belief over the cue position and the trial category by inverting the generative model it has learned. They then convert this belief into a response that minimizes the loss/reward function of the task as formalized by Bayesian decision theory (Figure 1C, middle row) To account for deviations from the ideal observer, this model can be extended by adding potential sources of approximations and biases in the decision making process (Rahnev and Denison, 2018; Ma, 2019). Biases can arise at the perceptual, categorical and/or the response stage. The two principal forms of variability are observation noise and approximate computations (Beck et al. (2012)). Next, we briefly describe each source of suboptimality as it arises during the transformation from sensory observation to behavioral response (Figure 1C, bottom row).

#### Observations

Observations are inherently noisy and possibly ambiguous giving rise to observation noise. Here, we include uncertainty from both external factors and internal factors, such as noise in the sensory periphery. In many cases, the magnitude of observation noise depends on the position of the cue. Such a dependence has been extensively studied in the context of Weber’s law, Steven’s power law etc. We follow Acerbi et al. (2014) who showed that such dependencies of observation noise on position can be accounted for by defining a nonlinear mapping from the external to an internal coordinate system in which noise is additive and Gaussian. For our later data analysis, we will fit this mapping directly to data (see Methods, and Figure 2 Figure supplement 1 for an illustration of this mapping).

#### Perceptual inference

We assume that during perception, the brain infers beliefs about latent variables *x*, given its sensory observations, *o*. The resulting belief, *p*(*x|o*), is the product of the likelihood, *p*(*o|x*), that characterizes the observation process and the brain’s prior expectations about *x*. For low-level sensory latent variables like position, this relationship of sensory latent variables to sensory observations is learned over a long time, and generally assumed to be close to optimal. We therefore model deviations from optimality in the inference process by allowing the mean (perceptual bias) and variance of the observer’s prior to differ from the (optimal) task-defined distributions. This choice is also justified by the fact that the distribution of observations usually deviate drastically between experiments and natural sensory inputs, implying a mismatch in an observer’s natural prior (obtained through lifelong learning) and the optimal prior for the task.

#### Categorical inference

Binary discrimination tasks introduce a binary variable, *D*, corresponding to the trial category that mediates the relationship between stimuli and correct responses, a relationship that we assume the observer has learned through instructions and task experience. For simplicity, and consistency with the task analyzed in the second part of this paper, we consider a simple localization task in which category *D* = 1 corresponds to locations to the right of the midline, *x* > 0, and category *D* = −1 corresponds to *x* < 0. However, any classification tasks in which the positions corresponding to each category i.e. *p*(*x|D* = −1) and *p*(*x|D* = +1) differ (e.g. different overlapping distributions as in (Drugowitsch et al., 2016)) are equally covered by this framework.

Importantly, the experimenter-defined distribution over *o* and the observer’s perceptual prior over *x* specify a distribution over the trial category. For instance, in the absence of a perceptual bias, and in an experiment in which *o* < 0 and *o* > 0 occur equally often, then the implied distribution over *D* is flat, i.e. *p*(*D* = −1) = *p*(*D* = +1) = 0.5 (Figure 2B). On the other hand, if the observer has a negative perceptual bias, then they, over the course of the experiment, will more often perceive the category to be 1, rather than +1. However, this implied distribution over the trial category may now be in conflict with the experimenter’s feedback after each trial, which will typically (by design in most experiments) imply a balance of both categories (the case we assumed here). As a result, the observer may learn an additional bias over the categories to act in such a way as to (partially) compensate for their perceptual bias. In the generative model, such a bias is formalized as a categorical prior, which may deviate from both the distribution over *D* implied by the sensory prior, but also from the one implied by the relative frequency of correct choices of either kind (Figure 2E). We call this additional bias ‘categorical bias’ since it leads to the observer’s perceptual inference to be non-calibrated in the sense that its expectations about the observations now deviate from the actually observed distribution. In general, perceptual and categorical bias can act independently of each other and separating them using data may be a challenge, as shown in 2.1.1.

**Figure 2:**
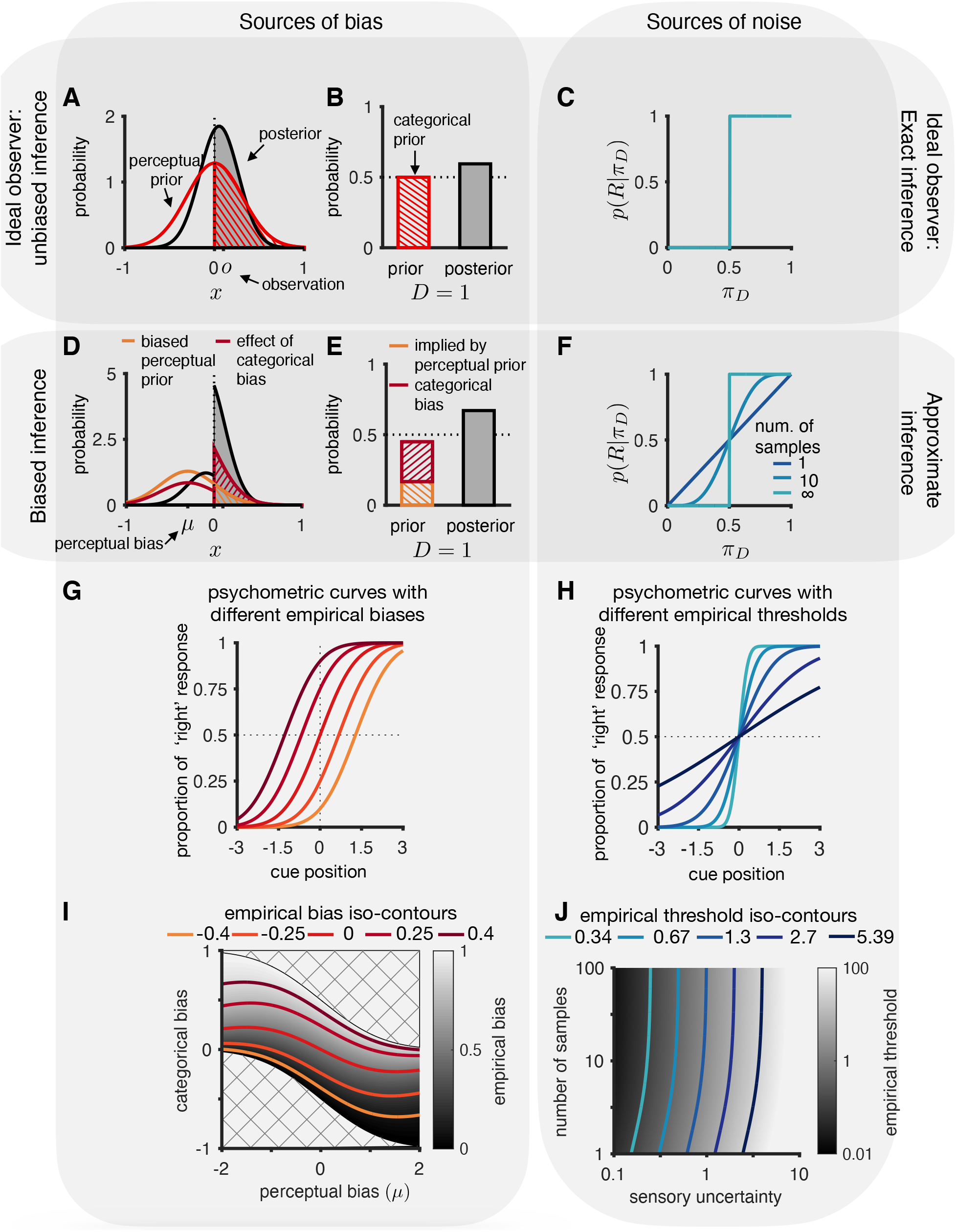
**(A-C)** Ideal observer for canonical task: zero mean prior that corresponds to the experimenter’s prior **(A)**, and 50-50 categorical prior **(B)** which we assume to be the correct one for the modeled experiment. The response is chosen that corresponds to the category with the highest posterior belief **(C)**. **(D-E)** Biased observer: a perceptual bias can be modeled as a prior whose mean is shifted away from zero (**D**, orange). A categorical bias manifests itself in a prior over the category *D* that differs from the distribution implied by the perceptual prior and the distribution over observations (**E**, red). **(F)** We model approximate computations by basing the response, *R*, on a finite number of samples drawn from the exact posterior, *π_D_*. One sample results in probability matching, and ∞ samples correspond to exact inference. **(G,I)** Any empirical bias can arise from different combinations of calibration and perceptual biases as illustrated by iso-contour lines in **(I)** for different empirical biases (corresponding psychometric curves shown in **G**). Hatched region indicates impossible combinations of perceptual and categorical bias. **(H,J)** As in **(G,I)** the observed empirical threshold can arise from different combinations of number of samples and observation noise as illustrated by iso-contour lines in **J** for different empirical log thresholds (corresponding psychometric curves shown in **H**)

#### Approximate computations and behavioral responses

Given a posterior belief over the trial category, the optimal response minimizes the expected task defined loss under the posterior. In the case of binary discrimination tasks with equal reward for either choice, this results in a strategy where it is optimal to report the trial category for which the posterior is highest (Figure 2C). The fact that inferences in the brain are approximate introduces yet another source of suboptimality. Since these approximations influence behavior by way of the categorical belief, *π_D_* = *p*(*D|o*), we model the aggregate effect of all computational approximations as resulting in an approximate posterior belief over the trial category. Inspired by sampling-based models of perception and cognition (Fiser et al. 2010), we chose to quantify the degree of approximation as the number of samples the brain uses to approximate the posterior. The approximation becomes more accurate as the number of samples increases, with infinitely many samples being equivalent to exact inference. The observer chooses the response corresponding to the belief for which most samples were generated. For the case of a single sample, this results in probability matching, while larger numbers of samples can be equivalently interpreted as a softmax response strategy with a temperature parameter that is inversely related to the number of samples (Drugowitsch et al., 2016). We emphasize that this parameterization of the degree of approximation does not commit our model to a neural sampling-based implementation but is simply an intuitive and general way to quantify computational precision.

Finally, we note that we also account for lapses (which are outside our Bayesian observer model framework) in order to model real data. We model lapses as occurring independently of the decision making process as shown in the generative model in Figure 1C. The lapse parameters in our model encapsulate any influences on the decision not yet captured, like choice error, loss of attention etc. Any motor-related response biases are also encapsulated in the lapse parameters that we fit for each observer (for details see Methods).

##### 2.1.1 Non-identifiability of different sources of suboptimality using observer responses in a binary discrimination task

Having defined different sources of suboptimality raises the question of whether they can actually be independently constrained using empirical data. It turns out that data from a simple binary discrimination task cannot distinguish between (a) perceptual bias and categorical bias and (b) degree of approximation (number of samples) and observation noise. This is because these four quantities combine to determine the empirically measured bias and sensitivity in a way that can not be disentangled. We have shown (Methods) that even with the extra suboptimality parameters, the observer response is given by the classic family of psychometric curves

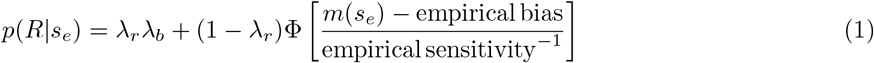

where *λ_r_* is the lapse rate, *λ_b_* is the lapse bias, Φ is the cumulative Gaussian distribution function, and *m*(*s_e_*) maps the external cue location onto internal coordinates to account for non-additive noise. Importantly, the empirical bias is a function of the perceptual bias, the categorical bias, the observation noise, and the prior variance. The empirical sensitivity is a function of the observation noise, the accuracy of the computational approximations, and the prior variance (see Methods for full analytical relationships).

#### Sources of bias

The empirical bias implied by the observer responses is the result of the priors in the *observer*’s generative model of the task differing from those used by the *experimenter*. The *ideal* observer’s priors over the cue position and the category match those of the experimenter. The prior over cue position is typically centered on zero and the prior over the category is typically 50-50 as illustrated in Figure 2A and 2B. When an observer has a biased perceptual prior (Figure 2D), this implies a categorical prior that differs from 50-50 (orange shading in Figure 2D and 2E). Furthermore, the observer’s categorical prior may be different from that implied by the perceptual prior, resulting in a categorical *bias* (Figure 2E indicated by the red shading). The categorical bias also scales the perceptual prior to reflect the different categorical prior than that implied by the perceptual prior (Figure 2D). An observer’s perceptual and categorical priors together determine its empirical bias. Equation (1) demonstrates that the two biases are indistinguishable given a single measured psychometric function. This is also illustrated by the iso-contours in Figure 2I where an observed empirical bias (Figure 2G) can arise from infinite combinations of perceptual and categorical biases.

#### Sources of noise

The observed empirical threshold depends on observation noise and the degree of computational approximation in the inference process. The ideal observer performs exact inference using the exact posterior probability over trial categories to choose the response that is most probable as illustrated in Figure 2C. Real observers, however, are necessarily approximate. We quantify the degree of approximation by the equivalent number of samples (infinite corresponding to exact inference, and one sample corresponding to probability matching). This results in a different response strategy as a function of the number of samples (Figure 2F). The observation noise and the number of samples together determine the empirical threshold. Equation (1) indicates that the two are indistinguishable given a single measured psychometric function. This is illustrated by the iso-contours in Figure 2J where an observed empirical threshold (Figure 2H) can arise from different combinations of observation noise and number of samples.

### 2.2 Choice irrelevant cues in a multi-sensory causal inference task can be used to dissociate different sources of suboptimality

We recently presented a task for which we demonstrated the ideal observer has a qualitatively different behavior than an approximate observer, regardless of their observation (sensory) noise (Cappelloni et al., 2019). This means that, in principle, it should be possible to use data from this task to infer sensory noise and degree of approximation separately. We will also show that, for this task, sensory prior and calibration prior have different effects on psychometric curves, implying that these two sources of suboptimality could, in principle, be dissociated using the empirical data.

First, we briefly summarize the task from Cappelloni et al. (2019): Two brief (300 ms) auditory stimuli, a tone and noise, were presented at equal eccentricity on opposite sides of the midline (Figure 3A). The observer was asked to report on which side of the midline the tone appeared. Temporally paired with the auditory stimuli, two random visual shapes were presented on the screen at different positions depending on the condition. In the first “central” condition, the visual shapes were presented on the midline, at the center of the screen. In the “matched” condition, the two visual shapes were presented at the same locations as the two auditory signals, tone and noise. Importantly, the appearance of the visual cues were random and not paired in any way with tone and noise and hence contained no information about the correct choice (left/right) in both conditions. In the matched condition, however, the visual cues did contribute information about the location of the auditory stimuli. The ideal observer in this task performs inference over whether they are in the central or matched condition: in the central condition, the visual cues should be ignored and in the matched condition, the observed locations of the visual cues should be cue-combined with the locations of the auditory cues. This process can be formalized as “causal inference” (Körding et al., 2007; Shams and Beierholm, 2010) and is shown as a graphical model in Figure 3B where the variable *C* represents the *condition* from the experimenter’s perspective, or the *causal structure* from the brain’s perspective and determines whether auditory and visual signals are combined (for equations see Methods).

**Figure 3:**
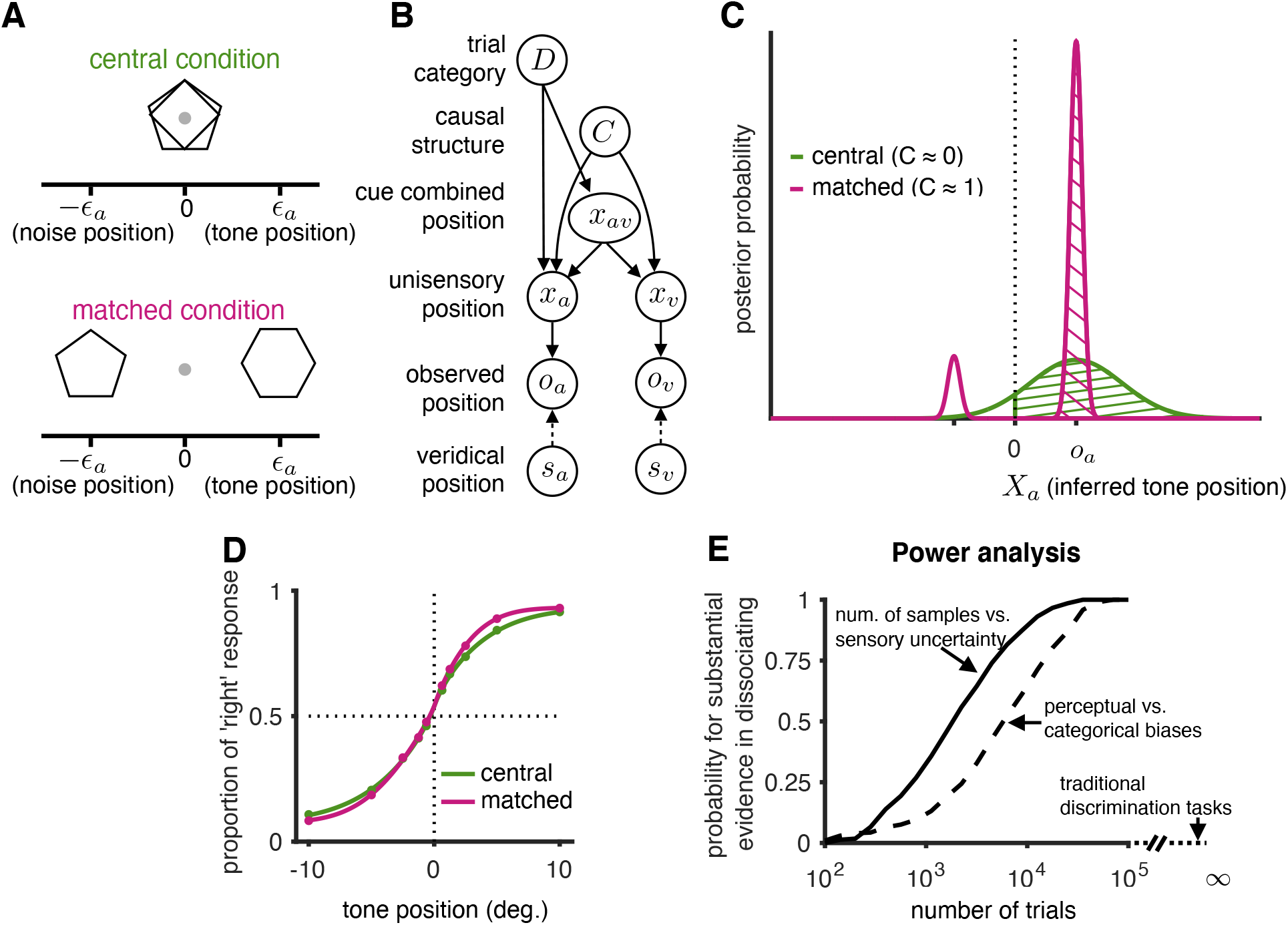
Task paradigm involving choice uninformative cues allowing for a dissociation of different sources of suboptimality. **(A)** On every trial, the observer observes four cues: a tone and a noise which are equidistant from the midline but on opposite sides and two visual cues (shapes) that are either overlapping at the midline in the central condition or aligned with the tone and noise in the matched condition. The pairing of shape and sound are random, making the visual cues uninformative about the correct choice. **(B)** Generative model for this task. The observer performs causal inference (Körding et al., 2007) to determine whether or not sounds and visual cues have the same eccentricity, and whether to combine information across them.**(C)** Illustration of how the visual cues affect the belief over the trial category. In the absence of visual cues, the belief over the trial category is the area of the likelihood function to the right of midline (assuming a flat prior over the inferred tone position and no biases). Assuming that the visual cues have a lower observation noise than the auditory cues, visual cues that are aligned (matched condition) will result in a bimodal likelihood function at the two cue positions. Since the larger of the two modes will always be on the same side as the mode of the auditory likelihood, the ideal observer will choose the same response in both conditions. However, since the left/right distribution of probability mass changes, an approximate observer (e.g. probability matching) may produce different responses in the two conditions **(D)** Predicted psychometric curves from an approximate Bayesian observer using realistic parameters (see fits to data below). **(E)** Power analysis that shows the probability of getting substantial evidence (measured using Bayes factor) in favor of two suboptimality extensions to the ideal observer model: approximate computations vs exact inference (solid line), categorical bias vs a perceptual bias (dashed line). Traditional binary discrimination task provide zero evidence in favor of both extensions. A similar plot using AIC for model comparison is shown in Figure 3 Figure supplement 1.

Importantly, an ideal observer, i.e. without any biases and performing exact inference, has the same psychometric functions in both the central and matched conditions. This is in line with expectations based on the fact that the visual cues by themselves contain no information about the correct choice. However, it turns out that the psychometric functions for each condition differ for an approximate observer in this task. Empirically the observer performance was also affected between central and matched conditions. Figure 3C recapitulates the visual proof (Cappelloni et al., 2019). In brief, in the central condition, when the auditory cues are not combined with the visual cues, the posterior may be a wide Gaussian centered on the observation. The ideal observer will report the side with the higher probability mass. However, in the condition in which the visual cues are matched to the auditory cues in eccentricity, the resulting posterior will be more highly localized around the eccentricities given by the visual cues. Importantly, while the side on which the probability mass is higher does not change leading an ideal observer to make the same decision in both conditions, the relative probability mass changes between conditions. As a result, the behavior of an approximate observer will be different since their ‘confusion probability’ will depend on the *relative* mass on both side. Note that the amount by which the approximate observer’s curves differ between the two conditions decreases with increasing approximation quality (parameterized by smaller number of samples in our case).

We performed a power analysis for the probability of finding substantial evidence in favor of approximate computations, where we defined ‘substantial’ as a Bayes factor greater than 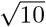 (Kass and Raftery, 1995) when compared against the exact inference model (Figure 3E). We simulated data from the causal inference model by choosing the sensory parameters as the average fit parameters across 20 observers for a particular value of number of samples and categorical bias (psychometric curve shown in Figure 3D). Given the number of trials available in our dataset (8000 trials across all participants), we had more power to constrain the computational approximation parameter independent of the sensory noise as compared to independently constraining sensory and categorical bias. We note that our estimate of the power is a conservative underestimate since the bias of the average observer considered here is smaller than that of the typical observer (since biases average out, unlike the computational approximation). We also performed the same power analyses using AIC instead of Bayes factor and obtained very similar results (see Figure 3 Figure supplement 1).

We next investigated the empirical signatures of the key suboptimalities in our model (Figure 4A). Traditional observation noise and perceptual bias, in the absence of computational approximations or a categorical bias, produce traditional sigmoidal psychometric curves that are identical for matched and central condition (Figure 4A “perceptual bias”). However, as soon as either the computations are approximate (Figure 4A “approximate inference), or the observer has a categorical bias (Figure 4A, “categorical bias”), the psychometric curves for central and matched condition deviate in characteristic ways. Despite the fact that the visual cues do not contain any information about the correct choice, an approximate observer’s performance improves substantially in the matched condition for intermediate eccentricities where the visual cues increase the certainty over the inferred category. A categorical bias, on the other hand, has an even more idiosyncratic effect on behavior. When it is “compensatory” to the perceptual bias, it reduces the overall bias as expected. However, it does so not by a simple shift in the psychometric function, but in a way that preserves the perceptual bias when the cue is at the discrimination boundary (zero), leading to a two-lobed adjustment to the psychometric curve (Figure 4A, “compensatory biases”). In our data, we find evidence for all these signatures of the different suboptimalities. Data from four example observers together with their model fits are shown in Figure 4B. We note that these signatures directly arise from the canonical extensions to the generative model, and were not the result of trying to post-hoc fit the data. In fact, it is hard to imagine an extension of a phenomenological model involving sigmoidal psychometric curves, e.g. by allowing for different slopes of biases across conditions, that would produce the predicted, and observed, behavior. (Also, see methods for a mathematical characterizations of these signatures.)

**Figure 4:**
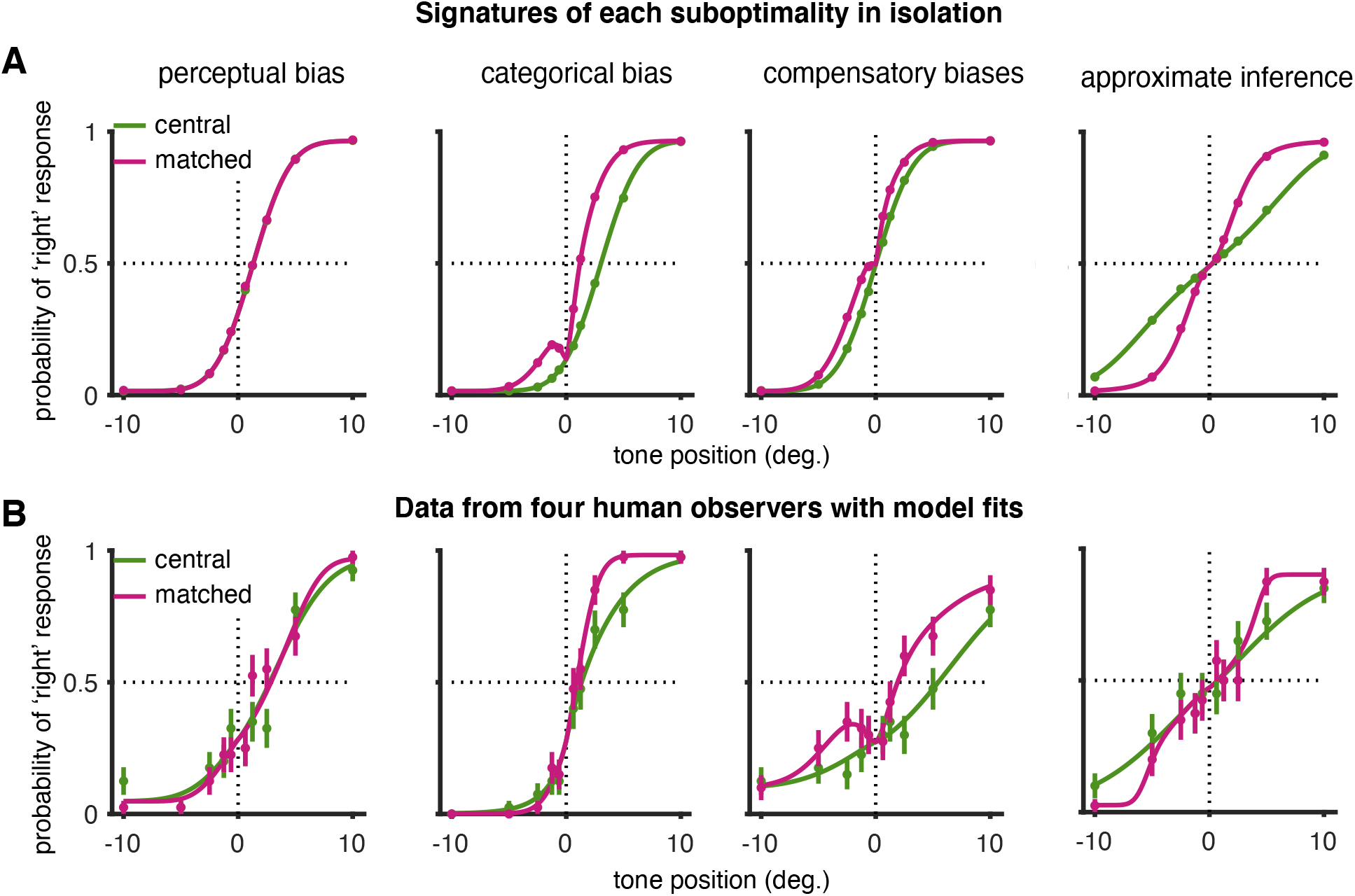
Empirical signatures of each suboptimality included in the model, and data from example observers. **(A)** Model predictions from left to right: (a) perceptual bias with no categorical bias and exact inference (b) categorical bias with no perceptual bias and exact inference (c) both perceptual and categorical biases such that the categorical bias is ‘perfectly’ compensatory, i.e. the empirically bias under this combination of perceptual and categorical biases is 0; exact inference (d) approximate inference with no bias. **(B)** Data from example observers responses and model fits: (a) perceptual bias with small categorical bias (b) categorical bias with small perceptual bias (c) both perceptual and categorical biases such that the categorical bias is compensatory, exact inference (d) approximate inference with no bias.

### 2.3 Analysis of behavioral data in choice-uninformative cue task

We next fit our approximate Bayesian observer model (Figure 3B) to the responses for each of the 20 individual observers in our data set (Cappelloni et al., 2019). We obtain full posteriors over all model parameters under weakly informative priors for each of the 20 observers (see Methods for fitting details). The key parameters of interest are the number of samples that quantify the degree of approximation and the categorical bias. Categorical bias represents the choice prior’s deviation from what is implied by the perceptual bias. The number of samples lies between one (coarsest approximation) and (exact inference). Since drawing 100 samples is indistinguishable from exact inference in our case, we only consider the range from 1 to 100, in addition to exact computations. We will compare the following three models: (a) exact inference without a categorical bias (but with a perceptual bias), (b) exact inference with perceptual and categorical bias, and (c) approximate inference with both kinds of biases and a finite number of samples characterizing the approximate computations. Approximate computations are the main focus for the remainder of the Results section.

Responses of two example observers were chosen to examine the model predictions more closely as shown in Figure 5A. Observer 1 was close to unbiased with approximate computations. Observer 2 suffered from both a perceptual and categorical bias and was also best explained assuming approximate computations. Data from both observers deviate from the assumption of exact inference (Figure 5A, dotted lines correspond to the null model, i.e. unbiased exact inference). All models explained the variance in the data reasonably well. There was a modest improvement for the model including suboptimalities, even when accounting for the increase in number of parameters (Figure 5 Figure supplement 1). We also performed a Bayesian model comparison using Bayes Factors (BF) to quantify how much one model is favored by the data compared to another model (Figure 5B). All our comparisons compare the full approximate inference model to its simpler alternatives, with BF’s larger than 1 indicating evidence in favor of the approximate model, and greater than 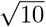 indicating substantial evidence in favor of the approximate inference model (Kass and Raftery, 1995). The data provides substantial support in favor of the approximate inference model over the exact inference model without categorical bias for ten of 20 observers and in favor of the approximate inference model over the exact inference model with categorical bias for 4 observers. Data from three of 20 observers provide strong evidence (BF> 10) for the approximate inference model as compared to the null model (unbiased exact inference).

**Figure 5:**
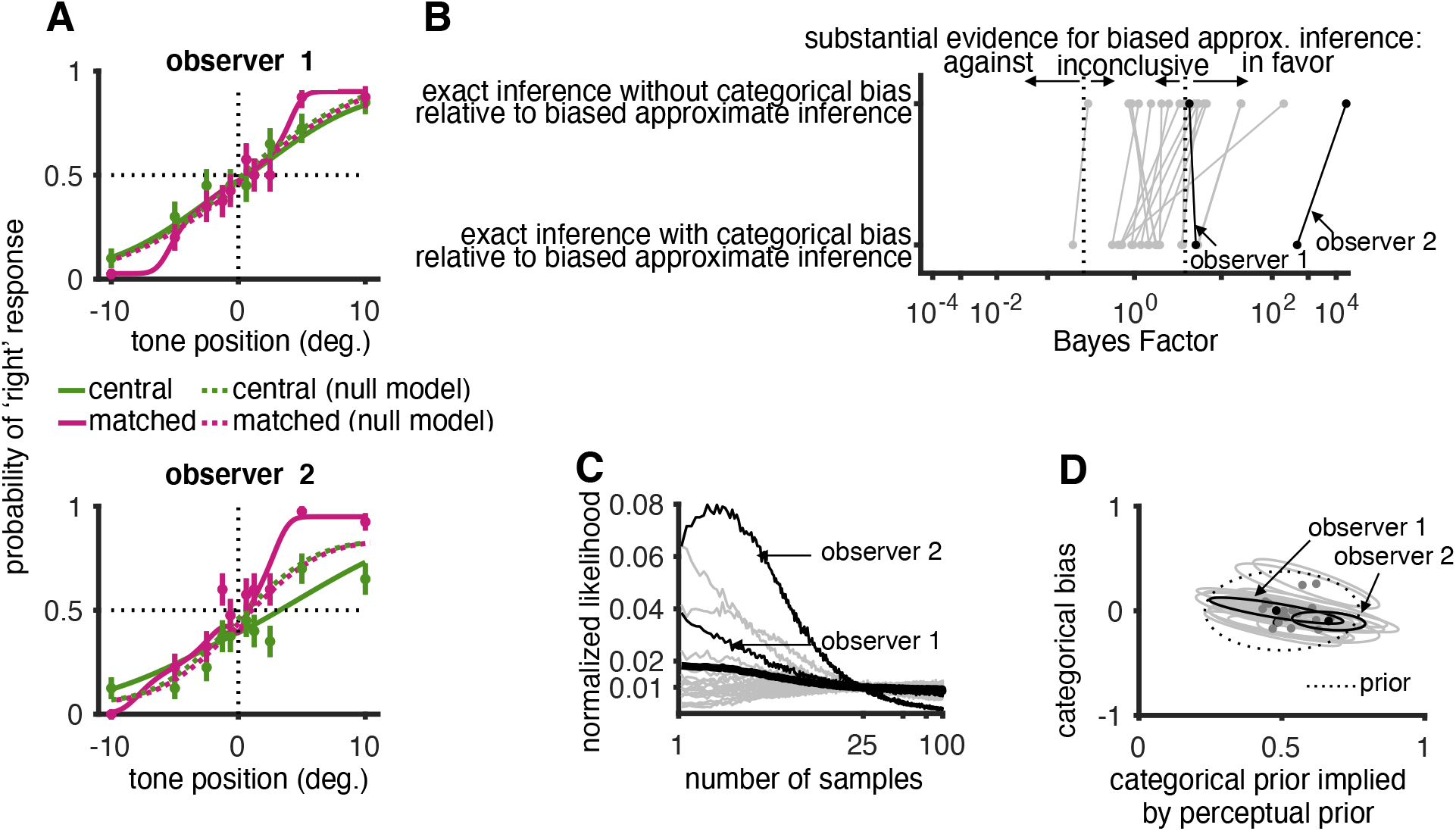
Model fitting and model comparison: **(A)** Responses from two example observers along with approximate Bayesian observer fits. Errorbars indicate 1 s.e.m. The dotted lines show the best fit ideal observer (unbiased and exact). **(B)** Model comparison between the approximate inference model and two alternate models: (i) exact inference model without categorical bias and (ii) exact inference with categorical bias. Positive Bayes factors indicate evidence in favor of the approximate inference model and values greater than 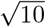 indicate substantial support for the approximate inference model. **(C)** Likelihood function showing likelihood of the data for different values of number of samples marginalizing out the other parameters. Thin graylines represents individual observers with the thin black lines indicating the example observers in A. The thick line represents average across observers. **(D)** Posterior distribution over the categorical bias for each observer indicated by thin gray lines with black lines indicating example observers in A.

The degree of approximation that best describes the data is shown individually for each observer in Figure 5C. Interestingly, for most observers, both extremes (one sample and 100 samples) are favored by the data, suggesting that there is a wide range of degrees of approximations underlying the task-relevant computations for the observers in our population. However, care must be taken to avoid over-interpreting the individual observer posteriors as they are wide due to the limited data per observer.

Figure 5D shows the joint posteriors over perceptual bias and categorical bias for individual observers. As expected from our power analysis (Figure 3E), for most observers there is little information constraining biases from the limited amount of data that we have for each observer. However, for a few observers (e.g. observer 2) the posteriors are markedly different from the prior clearly constraining both types of biases.

#### 2.3.1 Aggregate observer analysis

In order to pool statistical power across observers, we also perform our analysis on an aggregate observer whose responses are the combined responses across observers. The way we construct the aggregate observer from our data is equivalent to fitting a hierarchical model across observers allowing for variability across observers in all parameters except categorical bias and number of samples. Therefore the estimated categorical bias and number of samples are the average values across the population. We construct this aggregate observer by aligning the data from each individual observer in such a way as to account for their individual lapses, perceptual bias – all parameters except for number of samples and categorical bias (for details see Methods). If each observer performed exact inference and had no categorical bias, the transformed psychometric functions from all observers would be identical.

The aggregate observer data are shown in Figure 6A where the top panel shows the average psychometric curves for the central and matched conditions whereas the bottom row shows the average difference. The bottom panel is based on a paired comparison (difference) of matched and central condition per observer and clearly shows the hallmark of approximate inference in our task: enhanced performance for intermediate eccentricities, but little change for zero and large eccentricities. Our analysis of this aggregate observer strengthens the conclusion from the analysis of the individual observer data: the data decisively favors the approximate inference model (Figure 6C). The degree of approximation that best fits the data is based on a single sample, i.e. a coarse approximation. Its bias is not well-constrained by the data (Figure 6D). Finally, the full model accounts for marginally more variance in the data after accounting for the increase in number of parameters (Figure 5 Figure supplement 1).

**Figure 6:**
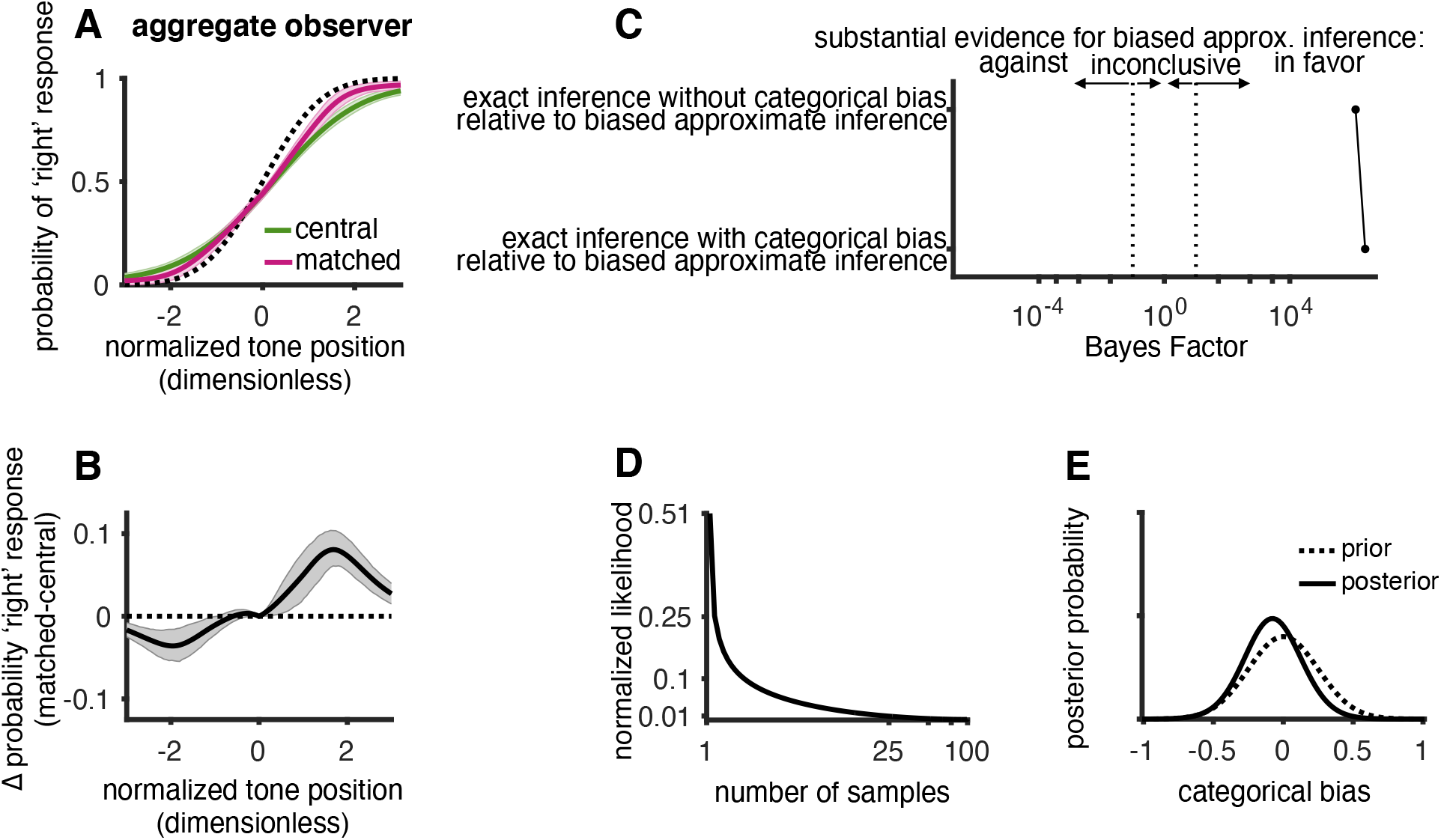
Model comparison for aggregate observer. **(A)** Model fits to aggregate observer data. Shading represents 1 s.e.m uncertainty intervals of posterior predictive distribution. The aggregate observer is constructed by combining normalized data from all individual observers after correcting for individual perceptual biases, observation noises and lapse parameters. The dotted line shows the fit ideal observer. **(B)** Mean difference between the probability of “right” responses between matched and central conditions condition. The dotted line shows the ideal observer prediction. Shading represents 1 s.e.m confidence intervals. **(C)** Model comparison (as in Figure 5C) shows overwhelming support for the approximate inference model over exact computations. **(D)** Likelihood of number of samples as in Figure 5D clearly favoring small number of samples. **(E)** The slight shift in the posterior over categorical bias for the aggregate observer provides weak evidence for a categorical bias on the population level.

### 2.4 Effect of stimulus duration on degree of approximation and sensory noise

Most theories of approximate inference (whether parameteric or sampling-based) predict that computations become more exact over time (Fiser et al., 2010; Pouget et al., 2013; Ma, 2019). However, testing whether this prediction holds in the case of the brain is complicated by the fact that observation noise is also likely to decrease with stimulus duration (Lengyel et al., 2015). Since our task allows us to dissociate the two of them, we repeated the experiment for different durations of the stimulus(Cappelloni et al., 2020). A change in sensory noise will manifest itself in a change in the overall slope of the psychometric functions. A change in computational accuracy affects both slope and the difference between the matched and the central conditions in a way that interacts with the observer’s biases.

Figure 7A shows the data from an example observer, showing a change in the difference between matched and central condition for increasing stimulus durations (top: 100 ms, middle: 300 ms, and bottom: 1000 ms). As expected, the overall slope of the psychometric functions increases with stimulus duration, compatible with a decrease in sensory observation noise. Additionally, the difference between the central and the matched conditions clearly decreases between 100 ms and 300 ms. While the change from 300 ms to 1000 ms is less obvious, model fitting confirms that the data for 300 ms favor a coarser approximation than the data for 1000 ms (Figure 7D).

**Figure 7:**
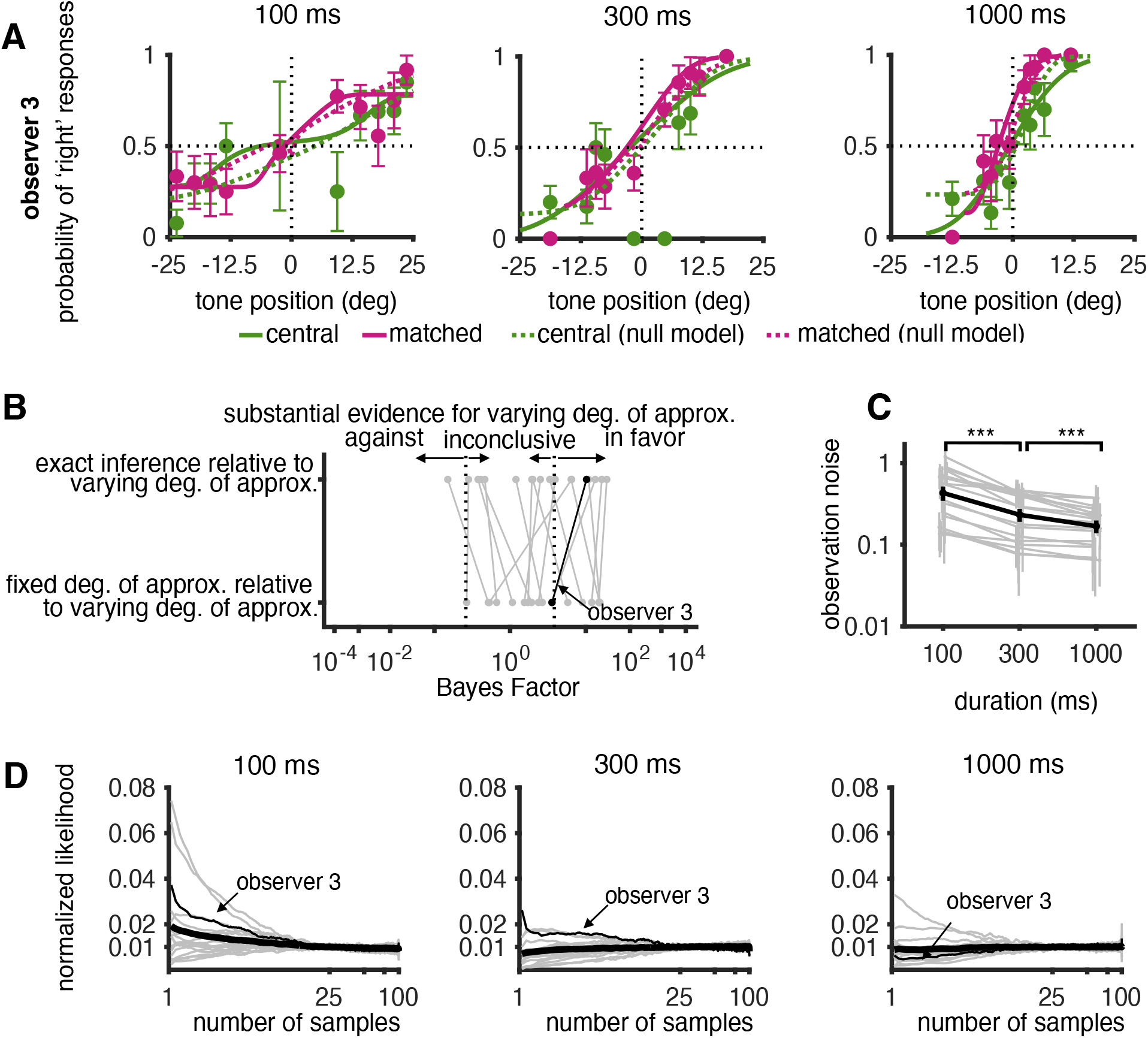
Individual observer analysis of variable duration task data: **(A)** Example observer psychometric curves (1 s.e.m. errorbars). Solid lines represent fits of approximate inference model whose degree of approximation varies with duration. Dotted lines represent exact inference model fits. **(B)** Model comparison of the approximate inference model with varying degree of approximation with two alternate models: exact inference model and approximate inference model with fixed degree of approximation. All models include perceptual and categorical biases. Positive Bayes factors indicate evidence in favor of the approximate inference model with varying degree of approximation with duration. **(C)** Observation noise as a function of duration for individual observers (thin gray lines) and population average (thick black line). Statistical significance assessed by a right tailed paired t-test (*p* = 2.5 × 10^−5^ for change from 100 ms to 300 ms and *p* = 2.8 × 10^−4^ for difference between 300 ms and 1000 ms). Significance is consistent under a non-parametric sign test. **(D)** Likelihood function for number of samples marginalizing out the other parameters for each duration. Individual observers (thin lines) and population average (thick line).

We perform Bayesian model comparison using Bayes Factors as before, now comparing the full approximate inference model with a degree of approximation (number of samples) that depends on stimulus duration, to two simpler models: first, (a) an approximate inference model with a fixed degree of approximation across all stimulus durations, and second, (b) an exact inference model (Figure 7B). Each of these three models includes for both perceptual and categorical biases. All models explained the variance in the data reasonably well (Figure 5 Figure supplement 1). We find that the data for eight of 20 observers provides substantial evidence in favor of approximate inference over exact inference, and for six of twenty observers in favor of a degree of approximation that changes with stimulus duration (Figure 7B).

Furthermore, and independent of our finding of the change in degree of approximation, we find that observation noise decreases as duration increases across the entire range of stimulus durations up to 1000ms (Figure 7C). As before, we find that the data from individual observers tends to favor either low or high numbers of samples, with a weak trend favoring higher numbers of samples for longer stimulus durations (Figure 7D). However, we clearly do not have enough data to draw reliable conclusions for most observers.

In order to again increase statistical power, we pool the data to form an aggregate observer (Methods). For all stimulus durations we find the pattern in the aggregate data that is characteristic of approximate inference (Figure 8A). Furthermore, we find that on the population level the data contains substantial evidence for an approximate inference model whose degree of accuracy changes with stimulus duration over an exact inference model and one whose degree of approximation is independent of duration (Figure 8B). Importantly, and in line with our initial hypothesis, we find that the data implies a systematic change from coarser to finer approximate computations as duration increases (Figure 8C).

**Figure 8:**
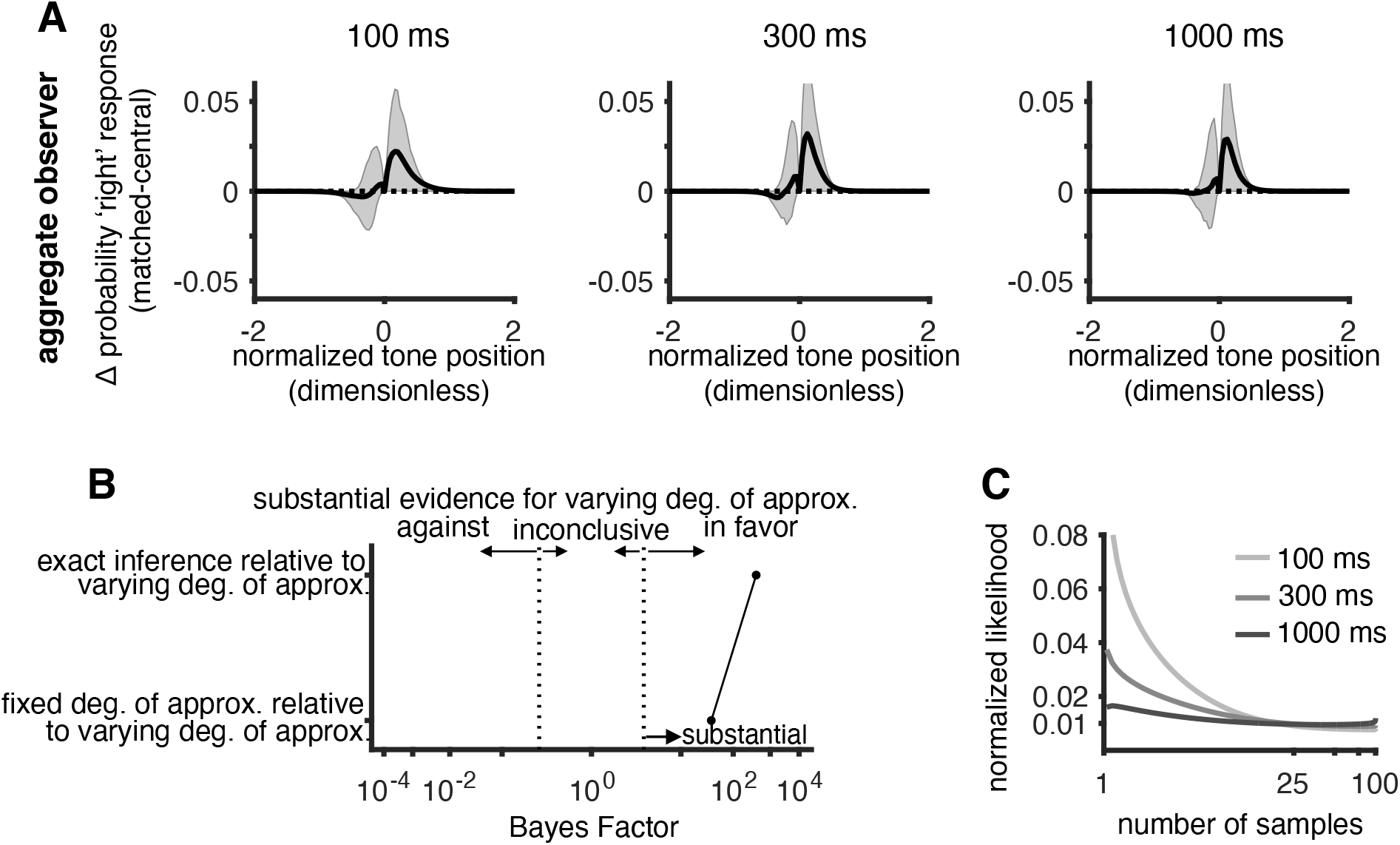
Aggregate observer analysis of variable duration task data:**(A)** Mean difference between probability of “right” responses in matched and central conditions (with 1 s.e.m. errorbars) where the aggregate observer is constructed analogously to Figure 6. The dotted line shows the prediction of the ideal observer (exact inference without categorical bias). **(B)** Model comparison between approximate inference model with varying degree of approximation and (i) exact inference and (ii) fixed degree of approximation. The aggregate data provides strong evidence that the degree of approximations depends on stimulus duration. **(C)** Likelihood functions for the number of samples. Change in shape reflects changing evidence for a coarse approximation for the shortest duration, to a better approximation for longer durations.

## 3 Discussion

Our work makes several conceptual and empirical contributions. First, we have extended the classic ideal observer model by four key sources of potential suboptimality in the context of the *generative* model, independent of how the inference process is implemented algorithmically. We showed that data from a simple discrimination task cannot dissociate the two sources of bias in our model, nor the two sources of variability (noise). Second, we showed that a more complex task involving causal inference containing choice irrelevant cues that affect the performance of suboptimal observers differently from ideal observers can dissociate between all these sources of suboptimality. Third, we used psychophysical data from that task to infer the sources of suboptimality for each observer. We found clear evidence for approximate computations, separate from observation noise, in both individual observers and on the population level. Finally, we found that both observation noise, and the accuracy of the approximate computations, improved with the duration of the presented stimulus across the entire range of tested stimulus durations: from 100 ms to 1000 ms.

Our Bayesian observer framework formalizes the prescription in (Rahnev and Denison, 2018) to construct observer models that quantitatively characterize the different sources of suboptimality in perceptual decision making. There are two principal ways in which suboptimality parameters can be added in a Bayesian framework: either by adding parameters to the components of the *generative* model (e.g. priors and likelihoods), or the components of the *discriminative* model (e.g. criterions or inference noise). Our work follows the the generative approach, in line with the idea that the brain learns a generative model of its inputs (“analysis by synthesis” Yuille and Kersten (2006); Lee and Mumford (2003)). Further it is related to earlier work quantifying deviations from optimality in a task by allowing for priors that were different from those defined by the experimenter (Stocker and Simoncelli, 2006; Odegaard et al., 2015; Noel et al., 2021). However, our approach differs from prior work that followed the discriminate model approach and added suboptimality parameters to the discriminative ideal observer model (e.g. (Brunton et al., 2013; Drugowitsch et al., 2016)). An important test for both generative and discriminative approaches will be how well the suboptimalities inferred in either framework will generalize to other tasks or contexts. As a practical concern we note that beyond extremely simple stimuli and tasks, the exact discriminative model quickly becomes complex and intractable, limiting the feasibility of the discriminative approach for natural or complex stimuli and tasks.

Taking a generative approach allows one to define the perceptual bias and the categorical bias in a way that directly relates to discrepancies between the statistics of natural inputs and those in a given task. Task statistics are rarely similar to natural statistics and earlier studies have indeed found that biased responses could be explained as a result of observers using their natural perceptual priors instead of those implied by a given experiment (Stocker and Simoncelli (2006); Odegaard et al. (2015)). How the brain resolves this conflict between priors learned outside and inside the context of a specific task might provide insights into the brain’s learning and compensation strategy among neurotypical and patient populations (see e.g. Noel et al. (2021)). By formulating decision priors as parameters in the generative model, we provide a systematic way of characterizing any deviation between decision and perceptual priors as categorical biases in the model, likely relevant for a wide range of studies into human decision-making.

Perceptual biases in decision making could also arise in the likelihood as opposed to the prior (Stocker and Simoncelli, 2005). While our formalization can easily be extended to allow for this additional source of suboptimality, under Gaussian assumptions for likelihood and prior, the biases in the likelihood and the prior are mathematically equivalent. Therefore, having an additional bias term in the likelihood would not change the explanatory power of the model in the context of our dataset.

We analytically showed the non-identifiability of the different sources of suboptimality. This was noticed earlier (Acuna et al., 2015; Linares et al., 2019; Wyart and Koechlin, 2016; Drugowitsch et al., 2016), leading to efforts to break them by using a more sophisticated task design. For example, Acuna et al. combined the results from an estimation task with those of a categorization task to dissociate sensory noise from approximate inference and did not find evidence of the latter. Drugowitsch et al. used an evidence integration task in which they varied the number of to-be-integrated stimulus frames in order to dissociate sensory from computational noise, finding that computational noise was an important source of behavioral suboptimality. Linares et al. combined data from two discrimination tasks with two different discrimination boundaries to dissociate between perceptual and categorical priors but did not distinguish between sensory noise and computational approximations. The data analyzed in our work has the advantage of being based on a single task (auditory discrimination with respect to the midline) with two randomly interleaved conditions (central and matched) of equal duration. This minimized changes between different tasks (e.g. estimation vs discrimination) potentially relying on different representations and decision strategies.

Our task design allows us to directly estimate the importance of observation noise and of approximate inference as a function of time. Traditionally, longer stimulus durations are thought to trigger an evidence integration process during which the brain averages away noise (Gold and Shadlen, 2007; Stine et al., 2020) explaining the observed improvement in behavioral performance. However, most approximate inference computations make the same prediction due to better approximations over time (Lengyel et al., 2015) – either due to their iterative nature as in MCMC sampling (Fiser et al., 2010) or stochastic implementation Pouget et al. (2013). Interestingly, we found evidence that both observation noise and degree of approximation changed across the entire range of stimulus durations tested (Figure 7C,8C;Figure 7 Figure supplement 1). Traditionally, perceptual inference has been conceptualized as a statistical problem of finding a signal in the noise (Swets et al., 1961). This framing has lead to non-bias sources of suboptimality being introduced as external and internal noise (Lu and Dosher, 2008). In our model, external noise is included in the observation noise while internal noise is split into two parts. Noise that is associated with the sensor generating the observation (e.g. the retina) is included in the observation noise. On the other hand, internal noise associated with increased variability during downstream computations contributes to approximate computations.

Our work also suggests a new explanation for improvements in observer performance in the presence of choice-uninformative cues in general: approximate inference. For instance, a previous study found that non-spatial auditory signals can improve performance during a visual search task in a cluttered environment (Van der Burg et al., 2008).

Our mathematical formalization also deviates from traditional approaches in two more minor ways. First, we allow for a flexible mapping from external to internal sensory “coordinates”. The observation noise in external coordinates has long been known to be stimulus dependent. However, the internal coordinates are chosen such that the observation noise in internal coordinates becomes additive and Gaussian, thereby making further inferences analytically tractable. This formulation is a modification of the mapping used by (Acerbi et al., 2014) providing a more intuitive understanding of the mapping parameters and contains purely linear (as used in most Bayesian models) and purely logarithmic (Fechner, 1860) as special cases instead of limiting cases as in (Acerbi et al., 2014). Second, we characterize lapses in observer responses using two parameters, lapse rate and lapse bias, reflecting an assumption about the underlying generative model about how they arise: as a fraction of trials in which the observer strategy is qualitatively different from their usual stimulus-based strategy, possibly reflecting lapses in attention as commonly assumed, or exploratory strategies (Pisupati et al., 2021).

Perceptual decision-making is suboptimal in many ways. How to best formalize the sources of these suboptimalities in order to arrive at models that better describe behavior, and that allow for deeper insights into the underlying beliefs and computations, is an open question (Rahnev and Denison, 2018). Our work demonstrates the usefulness of a formalization in terms of the generative model and approximate computations, when applied to data from a task of higher complexity than classic discrimination tasks.

## 4 Methods

### 4.1 Bayesian observer model for binary discrimination tasks

We model observer responses in a binary discrimination task using the Bayesian observer model presented in Figure 1C. The process of making a response based on sensory observations involves two stages: (a) the perceptual decision stage and (b) the response stage. Each stage describes a particular stage of the decision making process and is used to systematically parameterize any deviation from optimality. We use the case of auditory binary discrimination as an example for describing the model but the model is general for any task where the observer has to classify a presented cue between different categories. In the auditory discrimination task considered, the observer has to report which side of the midline, (i.e. a decision boundary), the auditory tone/cue came from.

We model the sensory observations of the observer as deviating from the veridical position due to external and internal observation noise. We model this as a zero mean additive noise added to the veridical position on each trial as given in Eq. (2)

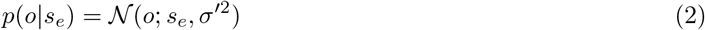

where 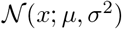 denotes the normal PDF with a mean *μ* and variance *σ*^2^. This model assumes that the observations are unbiased. The variance of the noise in general can be dependent on the tone position (Webers law, Stevens law etc), i.e. *σ*^′2^ = *f* (*s_e_*) where f is some function parameterizing the sensory noise. However, a position dependent variance makes the sensation distribution non gaussian which makes modeling further computations analytically intractable. However we can transform the cue position to internal coordinates (Acerbi et al. (2014)) such that the observation noise is cue independent in the internal coordinates as shown in Eq. (3)

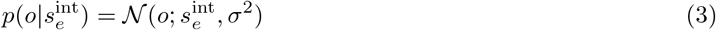

We use a transformation

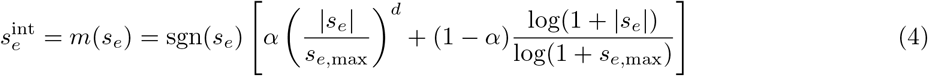

where *s_e_* is the veridical cue position and 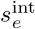 is the transformed value in internal coordinates. *α* controls the interpolation between a pure power law (Steven’s law, *α* = 1) to a purely logarithmic mapping (Weber’s law, *α* = 0). The exponent *d* models the power law coefficient where *d* = 1 corresponds to a purely linearly mapping which is the traditional assumption in Bayesian models for perceptual decision making. *s*_*e,*max_ is used to define the maximum value that the cue position can take such. This ensures any value when transformed to internal coordinates lies between −1 and 1. While the transformation holds for cue positions greater than *s*_*e,*max_, we can define the *s*_*e,*max_ as the edge of the screen in the experiment such that for any experiment, all cues lie within the specific maximum values. Since cue positions for most experiments defined in terms of visual angle are circular variables, the cue position in Eq. (4) can be shifted to lie within the principle range. The illustration of transformation of observation noise in internal coordinates is illustrated visually in Figure 2 Figure supplement 1

The perceptual decision stage describes how observers infer the beliefs about the latent causes that generated the sensory input. We consider two types of latent variables in the generative model: (a) perceptual latents that model the generative process for the observer’s observations and (b) task dependent latents that model the influence of task learning on the perceptual latents. In the discrimination task, the perceptual latent is the inferred tone position. We model all inferred variables to be in internal coordinates so we model the likelihood of the inferred tone position given in Eq. (5) to have the same form as Eq. (3) thereby assuming that the observers have learned a good estimate of their observation noise over lifelong learning.

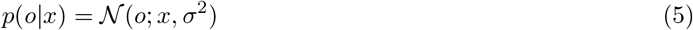

The task dependent latent is the decision variable (denoted by D) that represents the side of the decision boundary (assigned as the midline/zero without loss of generality) that the tone came from. The distribution over the inferred tone position conditioned on the decision variable is proportional to the product of two components as given in Eq. (6)

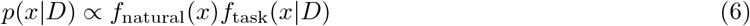

The proportionality constant can be obtained by integrating Eq. (6) over the support of *x*. The first component *f*_natural_(*x*) represents the natural prior over tone positions that the observer has learned over lifelong learning independent of performing the task. We model this component as a gaussian distribution as given in Eq. (7)

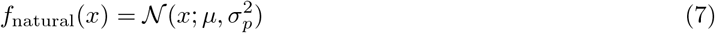

The second component partitions the space of *x* conditioned on the value of D. Denoting the value of D as 1 if the tone position is to the right of midline and −1 if the tone position is to the left of the midline, we define *f*_task_(*x|D*) as the approporiate partitioning of the space of *x* depending on D as written compactly in Eq. (8)

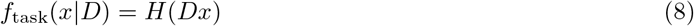

where *H*(*x*) is the heaviside function. The prior belief over D is modeled as a Bernoulli distribution with a prior 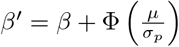 (Eq. (9)) where *β* is the categorical bias.

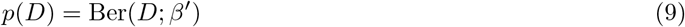

Defining the prior over *x* as given in Eq. (6) allows us to model task specific beliefs for each partition as specified by the prior belief over the decision variable (Eq. (9)) but also incorporate observer’s natural belief over the task variable (Eq. (7)) within each partition. The observer may or may not maintain separate task specific beliefs over the tone position. Our framework allows us to model the case where the observer does not have a separate task specific belief as is traditionally modeled when *β* is equal to the area of the natural component to the right of the midline which is 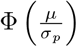 where Φ is the standard cumulative Gaussian distribution. Therefore any deviation of *β* from 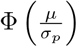 indicates the presence of a separate task specific belief that we refer to as categorical bias.

The response stage describes how observers convert their inferred belief about the decision variable into a response. Bayesian Decision theory provides a formal method of translating the posterior distribution over beliefs into observer responses by minimizing the task specific loss function, i.e.

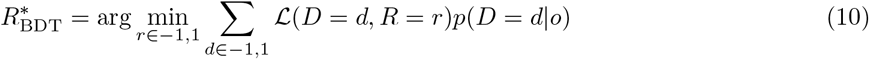

For the discrimination task with a 0-1 loss where the loss function is 0 if the observer reports the correct side and 1 otherwise, the optimal strategy is to choose 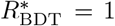 if *p*(*D* = 1|*o*) > 0.5 and 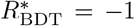 otherwise. However, the observer necessarily approximates the computation of *p*(*D* = 1|*o*) as exact Bayesian inference is intractable. We quantify the degree of approximation using a sampling based scheme where the observer uses a particular number of samples to approximate the posterior as given in Eq. (11)

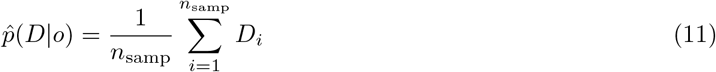

where *D_i_ ~ p*(*D|o*). Therefore under such an approximation scheme, the observer response strategy becomes

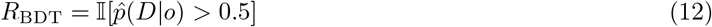

Intuitively, the observer draws *n*_samp_ samples from the posterior over D and chooses the side with the majority number of samples.

The observer responses could, however, deviate from *R*_BDT_ due to other external factors like attentional lapses, motor error etc. We model lapses as an independent corruption of the response (which we call lapse responses) by two variables lapse rate (*λ_r_*) and lapse bias (*λ_b_*). Lapse rate models the frequency of lapse responses and lapse bias models any bias towards D=1 while making a lapse response. This provides an alternate parameterization of two lapses that characterize the upper and lower offset in psychometric curve functions commonly used to model responses in binary discrimination tasks Fründ et al. (2011). Mathematically, this is given in Eq. (13)

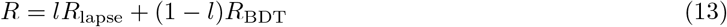

where *l* indicates whether or not lapse occured on a particular trial and is modeled as a bernoulli variable as given in Eq. (14) and *R*_lapse_ models whether the observer made a response 1 or −1 which is also modeled as a bernoulli variable as given in Eq. (15)

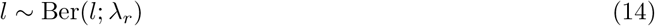

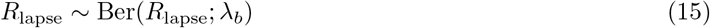

The distribution over observer responses for a given observation can be obtain by marginalizing over *l* and *R*_lapse_ in Eq. (13) using Eqs. (14) and (15) as given in (16)

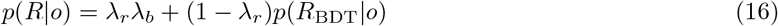

While Eq. (16) describes the observer response for a given observation, this cannot be measured directly by the experimenter. In turn they can only measure the observer response for a given *s_e_* which forms the psychometric curve. In order to evaluate the psychometric curve, we have to marginalize across all sensory observations in Eq.(16) as given in Eq. (17)

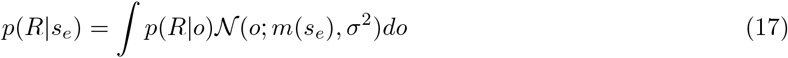

The integral in Eq.(17) is generally intractable but we derive an analytical approximation which is given in Eq. (18)

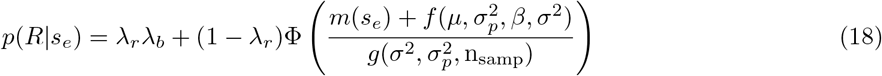

The effective bias 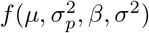 is

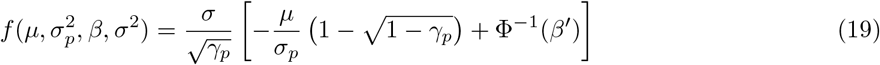

where *γ_p_* is the cue combined weight on the observation, i.e. 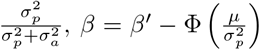 is the categorical bias and Φ^−1^ is the standard normal quantile function. The effective threshold is

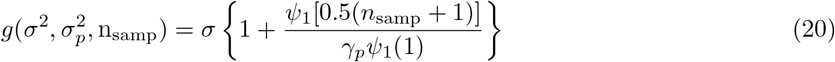

where *ψ*_1_ is the trigamma function. The full derivation of Eq. (18) is given in the following section.

### 4.2 Derivation of Bayesian observer responses in binary discrimination task

In this section, we derive the analytical approximation for a Bayesian observer in binary discrimination tasks, i.e. Eq. (18) in the previous section. We start with the response function (Eq. (12)) where the probability of making a “right” response is given by

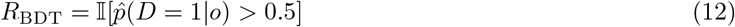

Given that 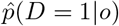 is a sample mean based estimate of *p*(*D* = 1|*o*) using *n*_samp_ samples, the probability that 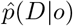 is greater than 0.5 is the probability that sum of *n*_samp_ random draws from the Bernoulli distribution with probability 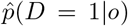 is greater than 0.5*n*_samp_. This can be written in terms of the Binomial CDF Φ*_b_* as given in Eq. (21)

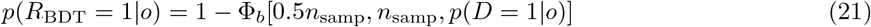

We can use the property relating the Binomial CDF to a Beta CDF to obtain a continuous expansion of Eq. (21) in terms of the beta cdf (Φ*_β_*) as given in Eq. (22)

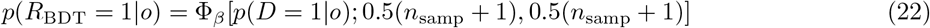

The advantage of Eq. (22) over Eq. (21) is that it provides an expression for the probability of the trial category that is approximated using *n*_samp_ and interpolates to continuous values of *n*_samp_. This is useful for optimization and analytic purposes. Eq. (22) can be rewritten in terms of the definition of the Beta CDF as given in Eq. (23)

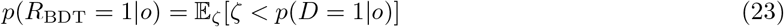

where *ζ* ∼ Beta[0.5(*n*_samp_ + 1), 0.5(*n*_samp_ + 1)] is a beta random variable. The inequality in Eq. (23) can be written in terms of the posterior odds over the trial category as shown in Eq. (24)

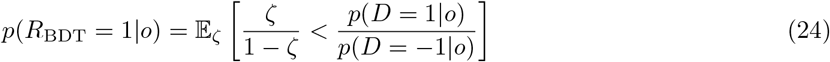

We can use Eqs. (5) to (9) to expand Eq. (24) to get Eq. (25)

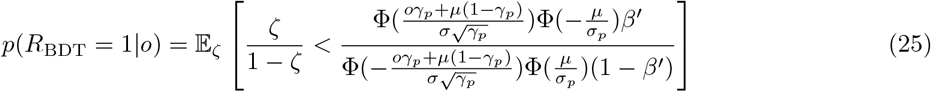

We can rearrange Eq. (25) as follows

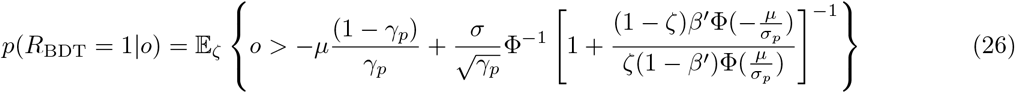

Eq. (26) shows an equivalence between the Bayesian observer and a signal detection theory model of decision making where the criterion is now stochastic for finite *n*_samp_ and reduces to a deterministic criterion for exact inference. Eq. (26) provides the probability of response for a particular sensory observation *o* which is inaccessible to the experimenter. The experimenter measures the probability of responding “right” for a value of *s_e_* that they vary to get the psychometric curve. Therefore, to get the probability of making a response *R_BDT_* = 1 for a given *s*, we have to compute the expected probability of Eq. (26) under the generative process of *o* given in Eq. (17) as shown in

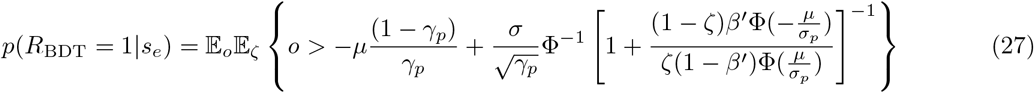

Since, the expectation is commutative, we can reorder the expectation above and write it in terms of the normal CDF to get

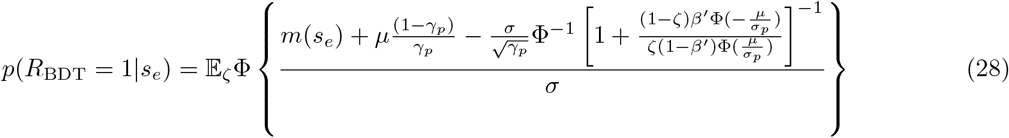

Using an approximation that Φ^−1^(*x*) ≈ *cs*^−1^(*x*) where *s* is the logistic sigmoid function (see equivalence analysis in (Drugowitsch et al., 2016)), we can simplify Eq. (28) to get

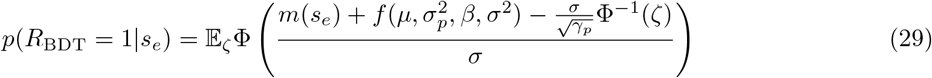

where

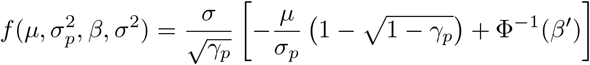

and 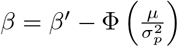 is the categorical bias

We can also approximate Φ^−1^(*ζ*) with moment matched gaussian distribution that has a mean of 0 and variance 2*c*^2^*ψ*_1_(0.5*n*_samp_ + 1) where *ψ*_1_ is the trigamma function and c is the approximation constant relating the cumulative normal cdf to a logistic sigmoid, i.e. Φ^−1^(*x*) ≈ *cs*^−1^(*x*). We can analytically evaluate the expectation in Eq. (29) under this normal approximation to get Eq. (18)

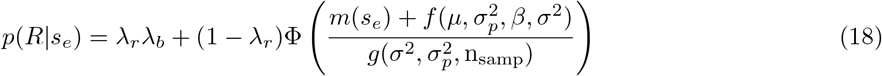

where

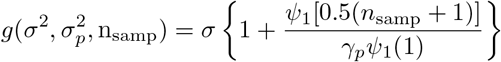

### 4.3 Task description for the choice uninformative cue tasks

We briefly describe the task in Cappelloni et al. (2019) which we refer to as the choice-uninformative cue task. On each trial, the observer observes two auditory stimuli: a tone (harmonics of 220 Hz) and noise. The noise was a randomly generated pink noise pattern with energy between 220 and 4000 Hz. Both the tone and noise had a with a 1/f spectral envelope. The auditory stimuli were accompanied with two visual shapes which were regular polygons inscribed in a circle with diameter 1.5 deg. The polygons were generated with constant luminance, saturation and opposite hues with number of sides randomly generated between four and eight such that the sides had different shapes on each trial. On every trial, the observer fixated at the center and observed the visual stimuli that appeared 100 ms before the auditory stimuli. The auditory stimuli were presented for 300 ms and both the auditory and visual cues ended together. The visual stimuli were presented at the midline in the “central” condition and aligned in eccentricity with the auditory stimuli in the “matched” condition. The observers had to report the side of the tone at the end of the trial. The eccentricities tested were 0.625, 1.25, 2.5, 5 and 10 degrees (both left and right) and there were 40 trials per condition resulting in a total of 800 trials across all conditions per observer.

In the variable duration version of the task (Cappelloni et al., 2020), the stimuli were the same but now the visual and auditory cue occurred concurrently. The four cues were presented for three different durations: 100 ms, 300 ms and 1000 ms. The stimuli were present in size 1 up 1 down staircase tracks that were interleaved with the separation between the tone and noise doubling when the observer made an incorrect response and reducing by a factor of 2^1/3^ when the observer made a correct response.

### 4.4 Bayesian observer model description for the choice uninformative cue task

We extend the Bayesian observer model presented in section 4.1 to model auditory discrimination in the presence of choice uninformative visual cues. The observer’s generative model in this task is given in Figure 3C where we present only the sensation and perception stage as the response stage is same as the Bayesian observer model in Figure 1C. The observer observes a tone and a noise which are drawn from the veridical values *s_a_* and *s_a_* corrupted by sensory noise as given in Eq. (30) where the cues are first transformed to internal coordinates using a mapping function *m* [Eq. (4)]

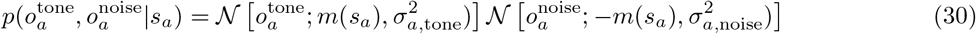

Similarly the observer observes a right and left visual cue which are drawn from the veridical values *s_v_* and —*s_v_* corrupted by sensory noise as given in Eq. (31) where the cues are first transformed to internal coordinates using a mapping function *m* [Eq. (4)]

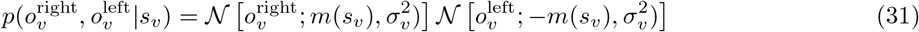

The likelihood of the inferred tone position mirrors the generative process as in the case of the Bayesian observer [Eq. (5)]. This is under the assumption that the observer has learned the task and combines information from both tone and noise observations to infer the belief over tone position as given in Eq. (32)

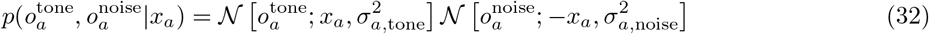

Similarly the likelihood of the right visual cue mirrors the generative process given in Eq. (31) under the assumption that the observer uses both right and left visual cue information to infer the belief over the right visual cue position as given in Eq. (33)

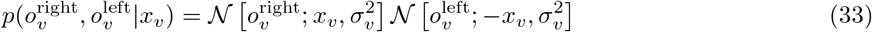

We model multisensory perception using a causal inference model where the observer infers whether or not the auditory and visual cues came from the same cause and use this to decide whether or not to combine information across the two cues. The model consists of four perceptual latents: (i) unisensory tone position (*x_a_*), (ii) unisensory right visual cue position (*x_v_*; right cue chosen without loss of generality), (iii) multisensory cue combined position (*x_av_*) and (iv) inferred causal structure (*C*). We also model task specific beliefs using a decision variable that indicates the side of the tone w.r.t the midline. The joint prior over the perceptual latents conditioned on the decision variable and the different values of the inferrred causal structure are given in (34) and (35)

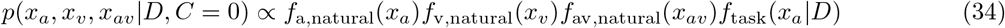

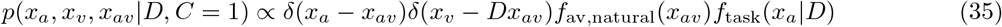

The proportionality constant can be obtained by integrating Eq. (6) over the support of *x_a_*,*x_v_* and *x_av_*. The first three components represents the natural prior over the tone, right visual cue and the auditory-visual combined cue positions which we model as a gaussian distribution as given in Eq. (36)

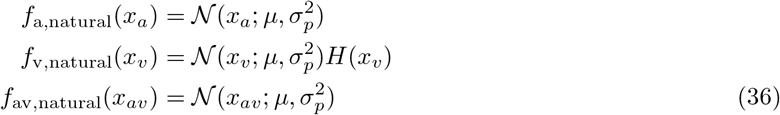

The fourth component models the influence of the decision variable on the perceptual latents as it partitions the space of *x_a_* conditioned on the value of D as given in Eq. (37)

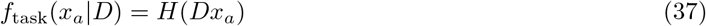

The prior belief over *C* is modeled as a Bernoulli distribution with a prior probability *p*_common_. Similarly, as was the case for the Bayesian observer, the prior belief over D is modeled as a bernoulli distribution with a parameter 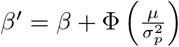. These two equations are given in Eq. (38).

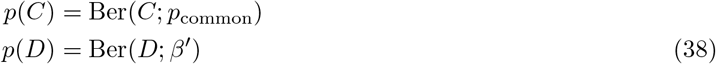

The mapping of the posterior probability over the decision variable to observer response is the same as the response stage described in section 4.1 [Eqs. (10) to (16)]. The observer response for given sensory observations can therefore be written as

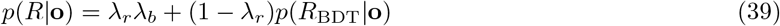

where 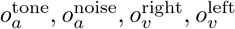 are abbreviated as **o**. *λ_r_* is the lapse rate which is the probability of making a lapse response and *λ_b_* is the lapse bias which is the probability of making a “right” response when lapsing. Further details about evaluating *p*(*R*_BDT_|**o**) are given in the. As in section 4.1, *R*_BDT_ is the response of the Bayesian observer approximated using *n*_samp_ samples

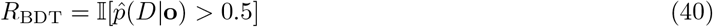

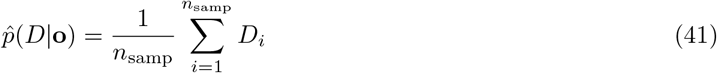

where *D_i_ ~ p*(*D*|**o**). In order to evaluate the psychometric curve for given experimenter defined positions, we have to marginalize across all sensory observations in Eq.(39) as given in Eq. (42)

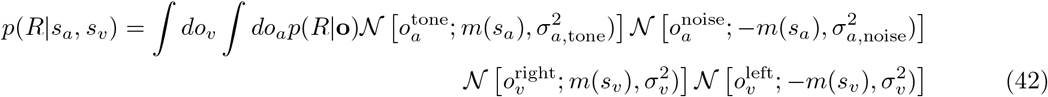

Since the integral in Eq. (42) is analytically intractable, we approximated the integrals using gaussian quadrature (Golub and Welsch, 1969). Gaussian quadratures provide a good approximation to the integral when *p*(*R*|**o**) is smooth. This is the case when *n*_samp_ is small. For larger values, the integrand becomes closer to a step function with the decision boundary becoming discontinuous. Therefore for the exact inference comparison models, we computed the decision boundary using a multi-dimensional bisection method (Bachrathy and Stépán, 2012) and then used this to compute the integral analytically. Given the predicted response from the model we can evaluate the likelihood of the observer responses **r** measured empirically for experimenter defined cue positions **s_a_**, **s_v_** as given in Eq. (43)

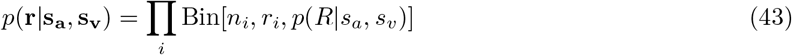

We obtained the maximum a posteriori (MAP) estimate for the model parameters under weakly informative priors using a quasi-newton Broyden-Fletcher-Goldfarb-Shanno (BFGS) unconstrained optimization procedure (fminunc in MATLAB) starting with 100 restarts to find the global optimum. We also obtained full posteriors over parameters using generalized elliptical slice sampling (Nishihara et al., 2014) which allowed us to get uncertainty estimates over parameter estimates.

In order get a goodness of fit estimate of the model, we computed the explainable variance explained (EVE) as described in (Haefner and Cumming, 2008). This estimate is an extension of the tradition variance explained/coefficient of determination but accounts for the uncertainty in the data generation process which is the case for us with limited number of trials per condition. In addition, it corrects for the number of parameters in the model to allow for overfitting. We also use Bayes Factor (Kass and Raftery, 1995) to perform model comparison. Computing the Bayes factor requires computing the marginal likelihood which requires evaluating the intractable integral in Eq. (44)

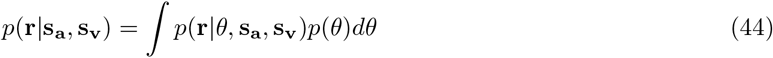

We approximate this using importance sampling as shown in Eq. (45)

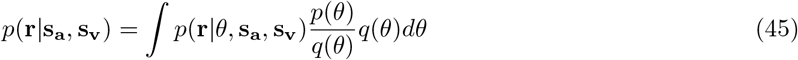

The quality of the approximation depends on how similar *q*(*θ*) is to the posterior *p*(*θ*|**r**, *s_a_, s_v_*). Therefore we approximate the posterior using a variational laplace approximation (Daunizeau, 2017) to approximate the posterior using a normal distribution and use that for evaluating the importance sampling based expectation. We improved the quality of the importance sampling approximation by using a Pareto fit smoothing to the weights as described in (Vehtari et al., 2015) and used a large number of samples (10000) to get a good estimate of the Bayes factor.

### 4.5 Derivation of observer responses in the choice uninformative cue task

We present a further derivation of the observer responses in the choice uninformative cue task and then present an approximation in the case when the visual uncertainty is much smaller than the auditory uncertainty. The response stage for the observer is the same as that described for the Bayesian observer in a binary discrimination task. Therefore the probability of response as predicted by Bayesian Decision Theory is

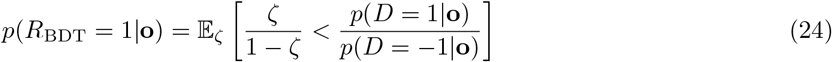

where 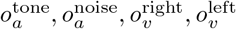 are together abbreviated as **o**. Since the experimenter only has access to *s_a_* and *s_v_*, in order to get the predicted probability of making a response *R_BDT_* = 1 for a given *s_a_* and *s_v_*, we have to compute the expected probability of Eq. (26) under the generative process of **o** given in Eqs. (30) and (31) as shown in

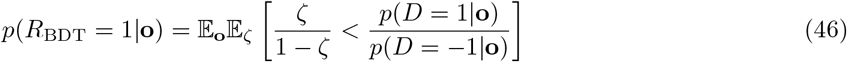

The belief about the trial category depends on the inferred causal structure and therefore we marginalize across the different causal structures to get *p*(*D* = 1|**o**)

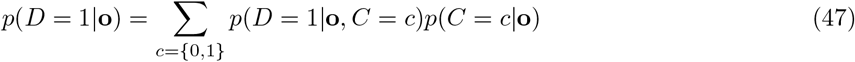

The conditional distribution *p*(*D* = 1|**o**, *C* = *c*) can be evaluated by marginalizing across the perceptual latents as given in

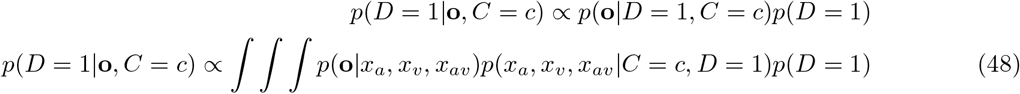

Using equations 23–29, we can infer the conditional belief over the trial category if the inferred causal structure is *C* = 0 as given in Eq. (49)

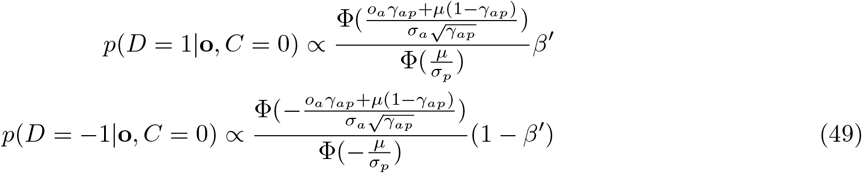

where 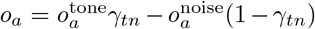 is the cue combined auditory position estimate, 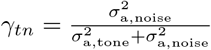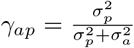 and 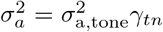. Similarly we can infer the conditional belief over the trial category if the inferred causal structure is *C* = 1 as given in Eq. (50)

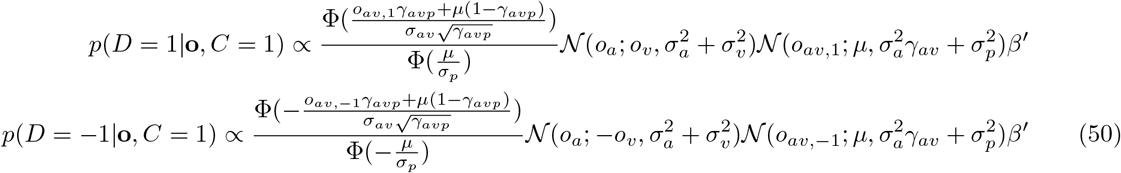

where 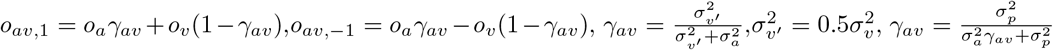. We can also evaluate the posterior probability over common cause using Eqs. (49) and (50) as given in (51)

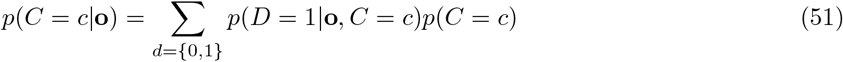

#### Approximate characterization of observer responses in the choice uninformative cue task

In order to get an interpretable functional form for observer responses, we make two assumptions: (a) For eccentricities sufficiently far from the midlines, the central condition always corresponds to the observer inferring *C* = 0 and the matched condition always corresponds to the observer inferring *C* = 1 (b) Also, we assume that the visual cue is very reliable, i.e. 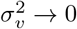

Substituting Eq.(49) in Eq.(46) and following the derivation similar to the traditional binary discrimination task, we can approximate the probability of observer response in the central condition as

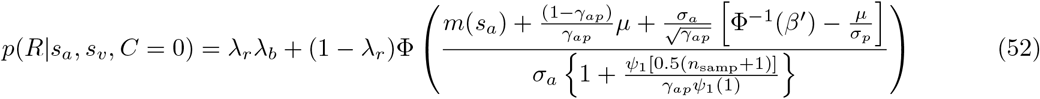

Similarly, the probability of observer response in the matched condition is given in Eq. (53)

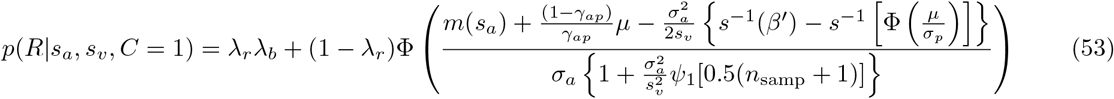

Intuitively, thresholds measured at different visual eccentricities from the matched condition allow us to separate the observation noise and number of samples. Measuring the biases for different visual eccentricities can allow us to separate the perceptual and categorical biases.

### 4.6 Combining observer responses into responses of an aggregate observer

While there is no analytical solution relating the observer responses to the experimenter defined cues positions in the choice uninformative cue task, we can approximate the response to a form (see previous section) as given below

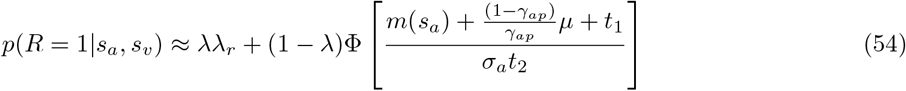

where *λ* is the lapse rate, *λ_r_* is the lapse bias, 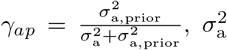 is the observation noise, *m*(*s*) is the mapping function and 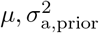 are the prior mean and variance over tone positions. The mapping function T maps experimeter defined locations *s_a_* to an internal measurement space such that any stimulus dependence on eccentricity is mapped to a stimulus independent uncertainty. *t*_1_ and *t*_2_ are functions of other parameters mainly visual cue location, categorical priors and number of samples respectively. *t*_1_ and *t*_2_ are 0 and 1 respectively, for observers who have no categorical biases (categorical prior matches perceptual prior) and perform exact inference. The left hand side of Eq. 1 is what we estimate by getting the proportion of rightward responses for a given *s_a_, s_v_* pair which we denote as *π_R_*. Since all the parameters differ from observer to observer, we can rewrite Eq. (54) for a observer *i* to get

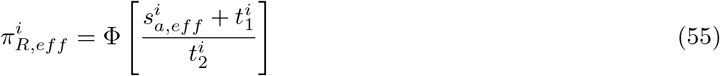

where 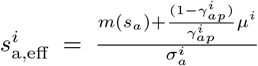 and 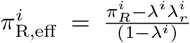. We can see from Eq. (55) that deviations of 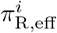 from 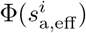 arise as a result of the observer having either a categorical bias or performing exact inference. Therefore if we combine responses across observers into an aggregate observer after mapping the 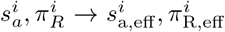, then any deviation between 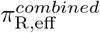 and 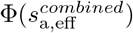 will contain information about the categorical biases and number of samples across observers. Since we do not have access to the true parameters for the observer, we sample from the posterior over the observer parameters. We therefore construct an aggregate observer using each parameter sample to get a distribution over aggregate observer responses. The number of trials that come from a observer for a given 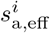 is (1 − *λ^i^*)*n^i^* where *n^i^* are the original number of trials for 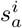 to compensate for the lapse responses made by the observer. Instead of just fitting *t*_1_ and *t*_2_ to the aggregate observer, we fit the full model to the aggregate observer to allow for deviations of sample estimate from the true value and the approximation involved in obtaining Eq. (54).

### 4.7 Modeling observer responses in the variable duration task

Unlike the fixed duration task where the observer responses were measured with a fixed set of stimuli, in the variable duration task, the observer responses were collected along a staircase procedure. The model in Figure 3C was fit separately to each duration while fixing the perceptual parameters across durations. In other words, we allow for different degree of approximation, sensory noise and lapse parameters for different durations while fixing the prior parameters. As in the previous section, we compare the approximate inference model to the exact inference model. We also compare the full approximate inference model to another approximate inference model where the degree of approximation is fixed across durations.

## Code and data availability

Code and data available at https://osf.io/6xbzt/

## Supplementary Figures

**Figure 2 Figure supplement 1**

Illustration of the cue-position dependent sensory noise obtained by using a non-linear transformation to internal coordinates and then adding a cue position independent sensory noise (Eq. (3)). The experimenter defined cue position, which we refer to as external position is transformed to internal coordinates with a mapping that lies between a linear and logarithmic mapping. If the mapping is linear, then the observation noise is independent of cue position in the internal coordinates. If the transformation is logarithmic, then the observation noise scales with cue position. This is depicted using the errorbars where the vertical errorbars indicate the observation noise in internal coordiantes that is cue independent. The horizontal errorbars depict the corresponding uncertainty in external position which scales with position for logarithmic and intermediate mappings

**Figure 3 Figure supplement 1**

Power analysis that shows the probability of getting substantial evidence (measured using AIC) in favor of two systematic extensions to the ideal observer model: solid line showing the approximate inference model as compared to a observer performance exact inference and dashed line showing the model having a categorical bias in addition to a perceptual bias as compared to a model having no categorical bias. Traditional binary discrimination task provide zero evidence in favor of both extensions.

**Figure 5 Figure supplement 1**

Absolute goodness of fit quantified by Explainable Variance Explained (EVE, (Haefner and Cumming, 2008)) which is the proportion of variance in the data that is predicted by the model adjusted for uncertainty in the data and number of parameters in the model. Goodness of fit presented for: **(A)** Fixed duration tasks for individual observers **(B)** Fixed duration tasks for aggregate observer **(C)** Variable duration tasks for individual observers **(D)** Variable duration tasks for aggregate observer

**Figure 7 Figure supplement 1**

Relative contribution of approximate inference to the measured threshold. For each observer, the relative threshold contribution is calculated as one minus the ratio of the threshold predicted under exact inference to the total threshold. The decrease in threshold contribution due to approximate inference is greatest for the shortest duration. Significance was assessed using a non parametric sign test and the decrease in threshold contribution from 100 ms to 300 ms was significant (*p* = 0.0013 for central and *p* = 0.001 for matched condition)

## Notes

### Competing Interest Statement

The authors have declared no competing interest.

### Summary of Updates

Figure 2 revised

