## Supplementary Information for "Inferring sources of suboptimality in perceptual decision making using a causal inference task"

#### Derivation of Bayesian observer responses in binary discrimination task

In this section, we derive the analytical approximation for a Bayesian observer in binary discrimination tasks, i.e. Eq. (18) in the main text. We start with the response function (Eq. (12)) where the probability of making a "right" response is given by

$$R_{\mathrm{BDT}} = \mathbb{I}[\hat{p}(D=1|o) > 0.5] \tag{12}$$

Given that $\hat{p}(D=1|o)$ is a sample mean based estimate of $p(D=1|o)$ using $n_{\mathrm{samp}}$ samples, the probability that $\hat{p}(D|o)$ is greater than 0.5 is the probability that sum of $n_{\mathrm{samp}}$ random draws from the Bernoulli distribution with probability $\hat{p}(D=1|o)$ is greater than $0.5n_{\mathrm{samp}}$. This can be written in terms of the Binomial CDF $\Phi_b$ as given in Eq. (39)

$$p(R_{\mathrm{BDT}}=1|o) = 1 - \Phi_b[0.5n_{\mathrm{samp}}, n_{\mathrm{samp}}, p(D=1|o)] \tag{39}$$

We can use the property relating the Binomial CDF to a Beta CDF to obtain a continuous expansion of Eq. (39) in terms of the beta cdf ($\Phi_\beta$) as given in Eq. (40)

$$p(R_{\mathrm{BDT}}=1|o) = \Phi_\beta[p(D=1|o); 0.5(n_{\mathrm{samp}}+1), 0.5(n_{\mathrm{samp}}+1)] \tag{40}$$

The advantage of Eq. (40) over Eq. (39) is that it provides an expression for the probability of the trial category that is approximated using $n_{\mathrm{samp}}$ and interpolates to continuous values of $n_{\mathrm{samp}}$. This is useful for optimization and analytic purposes. Eq. (40) can be rewritten in terms of the definition of the Beta CDF as given in Eq. (41)

$$p(R_{\mathrm{BDT}}=1|o) = \mathbb{E}_\zeta[\zeta < p(D=1|o)] \tag{41}$$

where $\zeta \sim \mathrm{Beta}[0.5(n_{\mathrm{samp}}+1), 0.5(n_{\mathrm{samp}}+1)]$ is a beta random variable. The inequality in Eq. (41) can be written in terms of the posterior odds over the trial category as shown in Eq. (42)

$$p(R_{\mathrm{BDT}}=1|o) = \mathbb{E}_\zeta\left[\frac{\zeta}{1-\zeta} < \frac{p(D=1|o)}{p(D=-1|o)}\right] \tag{42}$$

We can use Eqs. (5) to (9) to expand Eq. (42) to get Eq. (43)

$$p(R_{\mathrm{BDT}}=1|o) = \mathbb{E}_\zeta\left[\frac{\zeta}{1-\zeta} < \frac{\Phi(\frac{o\gamma_p+\mu(1-\gamma_p)}{\sigma\sqrt{\gamma_p}})\Phi(-\frac{\mu}{\sigma_p})\beta'}{\Phi(-\frac{o\gamma_p+\mu(1-\gamma_p)}{\sigma\sqrt{\gamma_p}})\Phi(\frac{\mu}{\sigma_p})(1-\beta')}\right] \tag{43}$$

We can rearrange Eq. (43) as follows

$$p(R_{\mathrm{BDT}}=1|o) = \mathbb{E}_\zeta\left\{o > -\mu\frac{(1-\gamma_p)}{\gamma_p} + \frac{\sigma}{\sqrt{\gamma_p}}\Phi^{-1}\left[1 + \frac{(1-\zeta)\beta'\Phi(-\frac{\mu}{\sigma_p})}{\zeta(1-\beta')\Phi(\frac{\mu}{\sigma_p})}\right]^{-1}\right\} \tag{44}$$

Eq. (44) shows an equivalence between the Bayesian observer and a signal detection theory model of decision making where the criterion is now stochastic for finite $n_{\mathrm{samp}}$ and reduces to a deterministic criterion for exact inference. Eq. (44) provides the probability of response for a particular sensory observation $o$ which is inaccessible to the experimenter. The experimenter measures the probability of responding "right" for a value of $s_e$ that they vary to get the psychometric curve. Therefore, to get the probability of making a response $R_{BDT}=1$ for a given $s$, we have to compute the expected probability of Eq. (44) under the generative process of $o$ given in Eq. (17) as shown in

$$p(R_{\mathrm{BDT}}=1|s_e) = \mathbb{E}_o\mathbb{E}_\zeta\left\{o > -\mu\frac{(1-\gamma_p)}{\gamma_p} + \frac{\sigma}{\sqrt{\gamma_p}}\Phi^{-1}\left[1 + \frac{(1-\zeta)\beta'\Phi(-\frac{\mu}{\sigma_p})}{\zeta(1-\beta')\Phi(\frac{\mu}{\sigma_p})}\right]^{-1}\right\} \tag{45}$$

Since, the expectation is commutative, we can reorder the expectation above and write it in terms of the normal CDF to get

$$p(R_{\text{BDT}} = 1|s_e) = \mathbb{E}_\zeta \Phi \left\{ \frac{m(s_e) + \mu\frac{(1-\gamma_p)}{\gamma_p} - \frac{\sigma}{\sqrt{\gamma_p}}\Phi^{-1}\left[1 + \frac{(1-\zeta)\beta'\Phi(-\frac{\mu}{\sigma_p})}{\zeta(1-\beta')\Phi(\frac{\mu}{\sigma_p})}\right]^{-1}}{\sigma} \right\} \tag{46}$$

Using an approximation that $\Phi^{-1}(x) \approx cs^{-1}(x)$ where $s$ is the logistic sigmoid function (see equivalence analysis in (Drugowitsch et al., 2016)), we can simplify Eq. (46) to get

$$p(R_{\text{BDT}} = 1|s_e) = \mathbb{E}_\zeta \Phi \left( \frac{m(s_e) + f(\mu, \sigma_p^2, \beta, \sigma^2) - \frac{\sigma}{\sqrt{\gamma_p}}\Phi^{-1}(\zeta)}{\sigma} \right) \tag{47}$$

where

$$f(\mu, \sigma_p^2, \beta, \sigma^2) = \frac{\sigma}{\sqrt{\gamma_p}} \left[ -\frac{\mu}{\sigma_p}\left(1 - \sqrt{1-\gamma_p}\right) + \Phi^{-1}(\beta') \right]$$

and $\beta = \beta' - \Phi\left(\frac{\mu}{\sigma_p^2}\right)$ is the categorical bias

We can also approximate $\Phi^{-1}(\zeta)$ with moment matched gaussian distribution that has a mean of 0 and variance $2c^2\psi_1(0.5n_{\text{samp}}+1)$ where $\psi_1$ is the trigamma function and c is the approximation constant relating the cumulative normal cdf to a logistic sigmoid, i.e. $\Phi^{-1}(x) \approx cs^{-1}(x)$. We can analytically evaluate the expectation in Eq. (47) under this normal approximation to get Eq. (18)

$$p(R|s_e) = \lambda_r\lambda_b + (1-\lambda_r)\Phi\left(\frac{m(s_e) + f(\mu, \sigma_p^2, \beta, \sigma^2)}{g(\sigma^2, \sigma_p^2, \text{n}_{\text{samp}})}\right) \tag{18}$$

where

$$g(\sigma^2, \sigma_p^2, \text{n}_{\text{samp}}) = \sigma\left\{1 + \frac{\psi_1[0.5(n_{\text{samp}}+1)]}{\gamma_p\psi_1(1)}\right\}$$

### Derivation of observer responses in the choice uninformative cue task

We present a further derivation of the observer responses in the choice uninformative cue task and then present an approximation in the case when the visual uncertainty is much smaller than the auditory uncertainty. The response stage for the observer is the same as that described for the Bayesian observer in a binary discrimination task. Therefore the probability of response as predicted by Bayesian Decision Theory is

$$p(R_{\text{BDT}} = 1|\mathbf{o}) = \mathbb{E}_\zeta\left[\frac{\zeta}{1-\zeta} < \frac{p(D=1|\mathbf{o})}{p(D=-1|\mathbf{o})}\right] \tag{42}$$

where $o_a^{\text{tone}}, o_a^{\text{noise}}, o_v^{\text{right}}, o_v^{\text{left}}$ are together abbreviated as $\mathbf{o}$. Since the experimenter only has access to $s_a$ and $s_v$, in order to get the predicted probability of making a response $R_{BDT} = 1$ for a given $s_a$ and $s_v$, we have to compute the expected probability of Eq. (44) under the generative process of $\mathbf{o}$ given in Eqs. (21) and (22) as shown in

$$p(R_{\text{BDT}} = 1|\mathbf{o}) = \mathbb{E}_\mathbf{o}\mathbb{E}_\zeta\left[\frac{\zeta}{1-\zeta} < \frac{p(D=1|\mathbf{o})}{p(D=-1|\mathbf{o})}\right] \tag{48}$$

The belief about the trial category depends on the inferred causal structure and therefore we marginalize across the different causal structures to get $p(D=1|\mathbf{o})$

$$p(D=1|\mathbf{o}) = \sum_{c=\{0,1\}} p(D=1|\mathbf{o}, C=c)p(C=c|\mathbf{o}) \tag{49}$$

The conditional distribution $p(D=1|\mathbf{o},C=c)$ can be evaluated by marginalizing across the perceptual latents as given in

$$p(D=1|\mathbf{o},C=c) \propto p(\mathbf{o}|D=1,C=c)p(D=1)$$
$$p(D=1|\mathbf{o},C=c) \propto \int\int\int p(\mathbf{o}|x_a,x_v,x_{av})p(x_a,x_v,x_{av}|C=c,D=1)p(D=1) \tag{50}$$

Using equations 23-29, we can infer the conditional belief over the trial category if the inferred causal structure is $C=0$ as given in Eq. (51)

$$p(D=1|\mathbf{o},C=0) \propto \frac{\Phi(\frac{o_a\gamma_{ap}+\mu(1-\gamma_{ap})}{\sigma_a\sqrt{\gamma_{ap}}})}{\Phi(\frac{\mu}{\sigma_p})}\beta'$$
$$p(D=-1|\mathbf{o},C=0) \propto \frac{\Phi(-\frac{o_a\gamma_{ap}+\mu(1-\gamma_{ap})}{\sigma_a\sqrt{\gamma_{ap}}})}{\Phi(-\frac{\mu}{\sigma_p})}(1-\beta') \tag{51}$$

where $o_a = o_a^{\text{tone}}\gamma_{tn} - o_a^{\text{noise}}(1-\gamma_{tn})$ is the cue combined auditory position estimate, $\gamma_{tn} = \frac{\sigma^2_{\text{a,noise}}}{\sigma^2_{\text{a,tone}}+\sigma^2_{\text{a,noise}}}$, $\gamma_{ap} = \frac{\sigma_p^2}{\sigma_p^2+\sigma_a^2}$ and $\sigma_a^2 = \sigma^2_{\text{a,tone}}\gamma_{tn}$. Similarly we can infer the conditional belief over the trial category if the inferred causal structure is $C=1$ as given in Eq. (52)

$$p(D=1|\mathbf{o},C=1) \propto \frac{\Phi(\frac{o_{av,1}\gamma_{avp}+\mu(1-\gamma_{avp})}{\sigma_{av}\sqrt{\gamma_{avp}}})}{\Phi(\frac{\mu}{\sigma_p})}\mathcal{N}(o_a;o_v,\sigma_a^2+\sigma_v^2)\mathcal{N}(o_{av,1};\mu,\sigma_a^2\gamma_{av}+\sigma_p^2)\beta'$$
$$p(D=-1|\mathbf{o},C=1) \propto \frac{\Phi(-\frac{o_{av,-1}\gamma_{avp}+\mu(1-\gamma_{avp})}{\sigma_{av}\sqrt{\gamma_{avp}}})}{\Phi(-\frac{\mu}{\sigma_p})}\mathcal{N}(o_a;-o_v,\sigma_a^2+\sigma_v^2)\mathcal{N}(o_{av,-1};\mu,\sigma_a^2\gamma_{av}+\sigma_p^2)\beta' \tag{52}$$

where $o_{av,1} = o_a\gamma_{av} + o_v(1-\gamma_{av})$,$o_{av,-1} = o_a\gamma_{av} - o_v(1-\gamma_{av})$, $\gamma_{av} = \frac{\sigma^2_{v'}}{\sigma^2_{v'}+\sigma^2_a}$,$\sigma^2_{v'} = 0.5\sigma_v^2$, $\gamma_{av} = \frac{\sigma_p^2}{\sigma_a^2\gamma_{av}+\sigma_p^2}$. We can also evaluate the posterior probability over common cause using Eqs. (51) and (52) as given in (53)

$$p(C=c|\mathbf{o}) = \sum_{d=\{0,1\}} p(D=1|\mathbf{o},C=c)p(C=c) \tag{53}$$

#### Approximate characterization of observer responses in the choice uninformative cue task

In order to get an interpretable functional form for observer responses, we make two assumptions: (a) For eccentricities sufficiently far from the midlines, the central condition always corresponds to the observer inferring $C=0$ and the matched condition always corresponds to the observer inferring $C=1$ (b) Also, we assume that the visual cue is very reliable, i.e. $\sigma_v^2 \to 0$

Substituting Eq.(51) in Eq.(48) and following the derivation similar to the traditional binary discrimination task, we can approximate the probability of observer response in the central condition as

$$p(R|s_a,s_v,C=0) = \lambda_r\lambda_b + (1-\lambda_r)\Phi\left(\frac{m(s_a)+\frac{(1-\gamma_{ap})}{\gamma_{ap}}\mu+\frac{\sigma_a}{\sqrt{\gamma_{ap}}}\left[\Phi^{-1}(\beta')-\frac{\mu}{\sigma_p}\right]}{\sigma_a\left\{1+\frac{\psi_1[0.5(n_{\text{samp}}+1)]}{\gamma_{ap}\psi_1(1)}\right\}}\right) \tag{54}$$

Similarly, the probability of observer response in the matched condition is given in Eq. (55)

$$p(R|s_a,s_v,C=1) = \lambda_r\lambda_b + (1-\lambda_r)\Phi\left(\frac{m(s_a)+\frac{(1-\gamma_{ap})}{\gamma_{ap}}\mu-\frac{\sigma_a^2}{2s_v}\left\{s^{-1}(\beta')-s^{-1}\left[\Phi\left(\frac{\mu}{\sigma_p}\right)\right]\right\}}{\sigma_a\left\{1+\frac{\sigma_a^2}{s_v^2}\psi_1[0.5(n_{\text{samp}}+1)]\right\}}\right) \tag{55}$$

Intuitively, thresholds measured at different visual eccentricities from the matched condition allow us to separate the observation noise and number of samples. Measuring the biases for different visual eccentricities can allow us to separate the perceptual and categorical biases.

#### Supplementary Figures

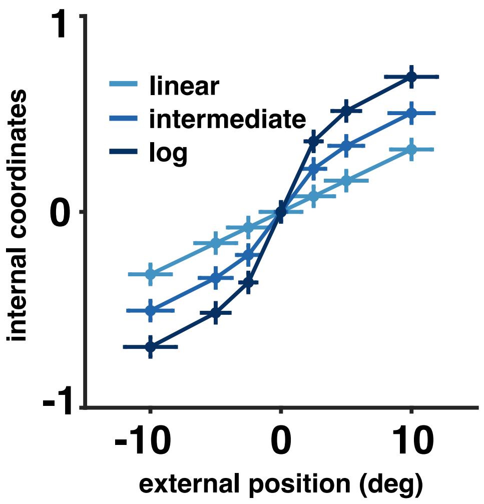


Figure S1: Illustration of the cue-position dependent sensory noise obtained by using a non-linear transformation to internal coordinates and then adding a cue position independent sensory noise (Eq. (3)). The experimenter defined cue position, which we refer to as external position is transformed to internal coordinates with a mapping that lies between a linear and logarithmic mapping. If the mapping is linear, then the observation noise is independent of cue position in the internal coordinates. If the transformation is logarithmic, then the observation noise scales with cue position. This is depicted using the errorbars where the vertical errorbars indicate the observation noise in internal coordiantes that is cue independent. The horizontal errorbars depict the corresponding uncertainty in external position which scales with position for logarithmic and intermediate mappings

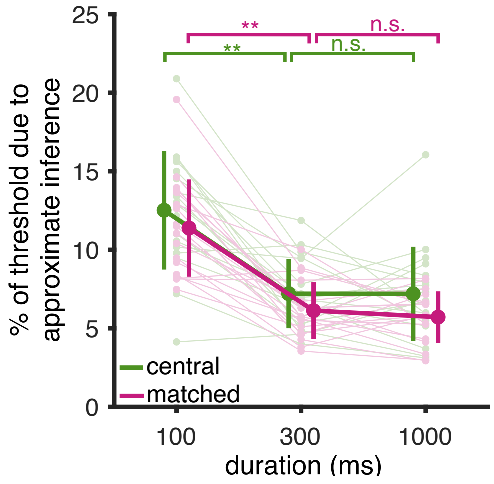


Figure S2: Relative contribution of approximate inference to the measured threshold. For each observer, the relative threshold contribution is calculated as one minus the ratio of the threshold predicted under exact inference to the total threshold. The decrease in threshold contribution due to approximate inference is greatest for the shortest duration. Significance was assessed using a non parametric sign test and the decrease in threshold contribution from 100 ms to 300 ms was significant ($p = 0.0013$ for central and $p = 0.001$ for matched condition)

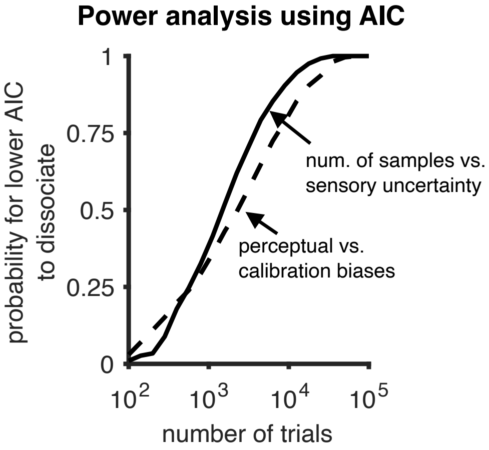


Figure S3: Power analysis that shows the probability of getting substantial evidence (measured using AIC) in favor of two systematic extensions to the ideal observer model: solid line showing the approximate inference model as compared to a observer performance exact inference and dashed line showing the model having a categorical bias in addition to a perceptual bias as compared to a model having no categorical bias. Traditional binary discrimination task provide zero evidence in favor of both extensions.

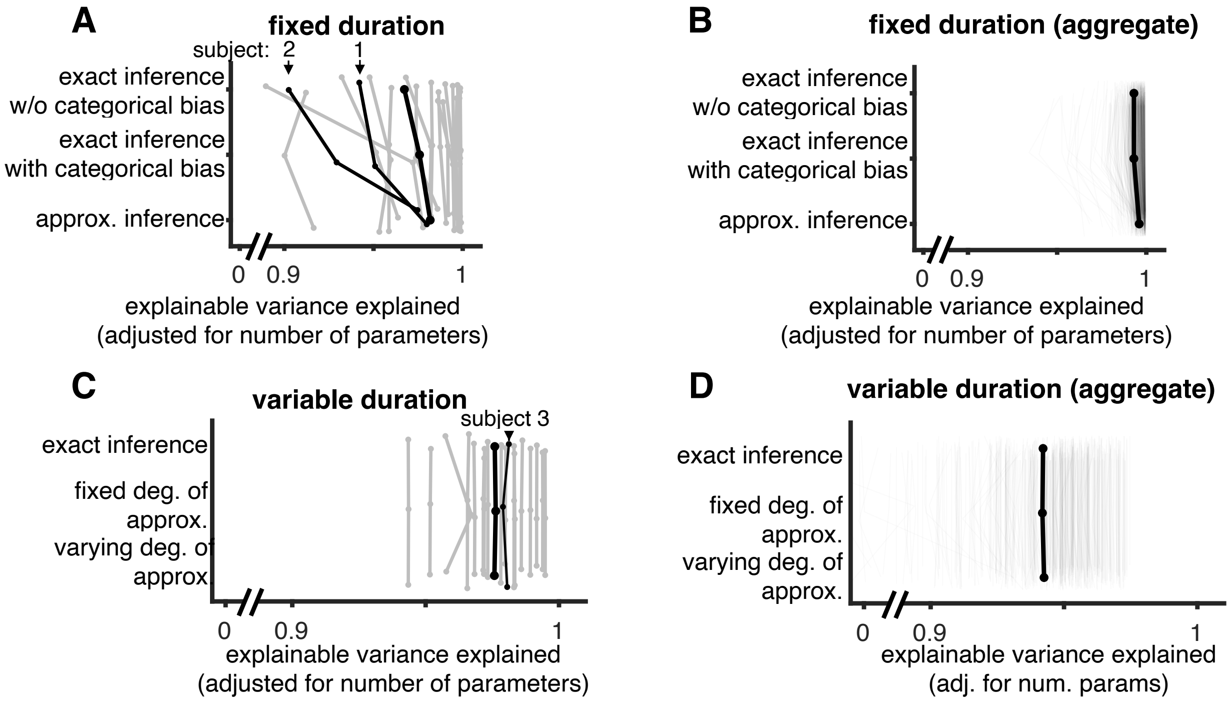


Figure S4: Absolute goodness of fit quantified by Explainable Variance Explained (EVE, (Haefner and Cumming, 2008)) which is the proportion of variance in the data that is predicted by the model adjusted for uncertainty in the data and number of parameters in the model. Goodness of fit presented for: **(A)** Fixed duration tasks for individual observers **(B)** Fixed duration tasks for aggregate observer **(C)** Variable duration tasks for individual observers **(D)** Variable duration tasks for aggregate observer
